# AI-Driven Design of Next-Generation Immunoinformatics Multi-Epitope Subunit Vaccine Targeting the Most Virulent Mpox Proteome: A Structural and Kinetic Validation Framework

**DOI:** 10.64898/2026.10.04.756542

**Authors:** Mohammed Emon, Noimul Hasan Siddiqque, Md. Enamul Haque

## Abstract

The resurgence of the Mpox virus (MPXV) highlights the urgent need for scalable, mutation-resistant countermeasures. Slow development timelines hinder traditional empirical vaccine discovery, while conventional vaccinology often yields high false-positive rates because it relies on static, linear sequence-based screening. These classical informatics filters consistently fail to predict physical structural stability or the dynamic presentation of immune cells under physiological conditions, creating a distinct translational gap between computational design and *in vitro* efficacy. To address these limitations, here we engineered an advanced, AI-augmented vaccine discovery pipeline targeting the most virulent components of the MPXV proteome. We systematically screened eight critical viral targets essential for replication, membrane morphogenesis, and host-cell entry to map highly conserved T-cell and B-cell epitopes. Moving beyond standard heuristic filtering, we implemented a high-fidelity deep learning validation layer: ESM-2 protein language models (pLMs) were deployed to calculate pseudo-perplexity scores across chimeric linker junctions (EAAAK, GPGPG) to guarantee structural naturalness, MHCflurry ensemble models successfully verified active intracellular antigen presentation, and structure-informed graph neural networks (GNNs) simulated the physical node-attention dynamics of the immune synapse. Finally, recurrent neural networks (RNNs)-driven mRNA kinetic modeling optimized translation initiation rates and minimum free energy profiles during *in silico* cloning. This multi-layered pipeline successfully yielded three structurally refined, multi-epitope vaccine constructs optimized for maximum global population coverage. ESM-2 structural validation confirmed exceptionally low perplexity scores at structural junctions, indicating that the engineered chimaeras have native-like stability and a minimized risk of proteolysis. GNN-based immune synapse modeling demonstrated highly stable binding kinetics and robust binding free energies between the target epitopes and human HLA alleles. Furthermore, RNN kinetic optimizations established that the cDNA sequences are optimized for high-yield expression in heterologous expression hosts without risk of ribosomal stalling or translational bottlenecks. This study establishes a deep learning-augmented framework that bridges the gap between theoretical sequence optimization and biopharmaceutical scalability. By validating the structural naturalness, real-world immunogenicity, and manufacturing kinetics of our vaccine chimeras, this framework presents a robust, proactive paradigm for rapid vaccine architecture against emerging orthopoxvirus threats.

**Key points:**

- A synthetic multi-epitope chimera (MPXV-3) was engineered across eight core MPXV virulence proteins utilizing rigid EAAAK, proteasomal AAY, and flexible GPGPG linkers to optimize structural spacing and processing.
- Screening and cross-validated candidate epitopes against mass-spectrometry-trained presentation models, confirming optimal intracellular processing and presentation ranks (0.5%) across prevalent HLA Class I alleles, led by FLADKKMTL (*K_D_* = 15.07 nM, percentile rank: 0.002%) and NLLCHIYSL (*K_D_* = 15.32 nM, percentile rank: 0.098%).
- The construct demonstrated dual-arm immune competence by engaging the innate TLR3 receptor in full-length protein docking, alongside structure-informed graph neural networks (GNNs) modeling of the ternary TCR–pMHC-I interface to verify high-affinity synapse formation (*P*_binder_ >0.70).
- Generative mRNA sequence modelling yielded a stable full-length transcript (ΔG = −678.1 kcal mol^-1^) paired with an unconstrained single-stranded 5’ initiation portal (ΔG = −11.6 kcal mol^-1^), preventing secondary structure blocks during ribosomal scanning.
- The codon-optimized coding sequence was computationally verified for insertion into the pET-28a(+) expression vector and *XhoI* restriction sites, maintaining reading-frame continuity with a C-terminal 6xHis-tag while confirming the absence of internal restriction cut sites.

## Introduction

The persistent threat of global viral resurgence, exemplified by the rapid escalation and multi-jurisdictional spread of the Mpox virus (MPXV), has exposed critical vulnerabilities in the global countermeasure infrastructure [1], [2]. While first- and second-generation orthopoxvirus vaccines provided essential baseline cross-protection during historical outbreaks, their current clinical utility in widespread populations is severely hampered by systemic adverse reactogenicity, complex manufacturing scaling, and a distinct lack of precise multi-antigen targeting designed to prevent mutational escape [3], [4].

Traditional empirical vaccine discovery, reliant on iterative *in vitro* screening and animal modeling, remains a prohibitively slow and resource-intensive paradigm that routinely fails to react at the speed of accelerating viral evolution [5], [6], [7]. To bypass these operational bottlenecks, reverse vaccinology and immunoinformatics have emerged as dominant methodologies for the rapid, genome-wide identification of candidate T-cell and B-cell epitopes [6], [8].

However, conventional *in silico* workflows frequently suffer from high false-positive rates because they rely almost exclusively on static, linear sequence-based prediction heuristics [9], [10]. These traditional matrix-based filters regularly fail to account for the highly dynamic, non-linear physical interactions and thermodynamic constraints that govern antigen presentation and structural conformation under physiological conditions [11].

Consequently, many computationally optimized vaccine chimeras suffer from local structural misfolding, rapid proteolytic degradation, or suboptimal surface presentation when translated from AI digital models to *in vitro* expression [12], [13]. To address these critical translational limitations, we upgraded our computational pipeline by integrating a high-fidelity AI-validation layer that utilizes graph neural network (GNNs) to simulate the physical dynamics of the immune synapse and ESM-2 protein language models (PLMs) to verify the structural stability of the multi-epitope vaccine chimera, while simultaneously implementing RNN-driven mRNA kinetic modeling to optimize translational efficiency and ensure the candidate’s viability for large-scale industrial manufacturing [14], [15].

By focusing specifically on the most virulent components of the MPXV proteome, we systematically selected eight critical targets essential for viral replication, membrane morphogenesis, and host-cell entry, namely, intracellular viral protein, viral membrane formation protein, virion protein D2, virion core protein D3, membrane protein CL5, assembly protein G7, virion core protein P4b, and viral core cysteine proteinase. This computational vaccinology study moves beyond simple and basic predictive screening to establish a robust and holistic deep learning-augmented framework that effectively bridges the gap between theoretical sequence optimization and biopharmaceutical scalability, presenting a robust and proactive strategy for targeted vaccine architecture against emerging orthopoxvirus threats [16], [17].

This study implemented an integrated, multi-tiered computational framework combining evolutionary protein language modeling, deep learning-augmented immunopeptidomics, and structural topological evaluation to identify and validate protective epitopes against the virulent Mpox virus (MPXV). Through systematic screening and filtering across diverse viral targets, the candidate construct MPX-3 emerged as our prioritized lead candidate. Driven by the core immunogenic nonamer FLADKKMTL, MPX-3 was selected based on its favorable intracellular processing profile, high-affinity human leukocyte antigen presentation (HLA-A*11:01), and structural stability across the simulated T-cell receptor (TCR)–pMHC immune synapse. By subjecting MPX-3 to rigorous latent-space perplexity assessment and graph neural network (GNN) synaptic topology profiling, this work provides a promising biophysically grounded blueprint for next-generation, structure-validated Mpox vaccine design.

## Methodology

### Strain identification, selection, and retrieval of the protein sequences

We selected Mpox virus strains by reviewing numerous entries in the NCBI online database. Eight protein sequences such as intracellular viral protein (accession no: NP_536598), viral membrane formation protein (accession no: NP_536549.1), virion protein D2 (accession no: NP_536526.1), virion core protein D3 (accession no: WQK80685.1), membrane protein CL5 (accession no: NP_536511.1), assembly protein G7 (accession no: NP_536504.1), virion core protein P4b (NCBI reference sequence: NP_536541.1) and viral core cysteine proteinase (GenBank: UVT84927.1) were downloaded from the NCBI database in FASTA file format.

### Allergenicity, toxicity, and physicochemical property analysis of the protein sequences

The antigenicity of protein sequences was determined using the online tool VaxiJen (v2.0) [18]. For antigenicity prediction, we set the accuracy threshold to 0.45 for the tumor model [19], [20]. We used the tumor model because it generated excellent results (compared with other Models) in both leave-one-out cross-validation (LOO-CV) and external validation in numerous studies. The accuracy, sensitivity, and specificity of VaxiJen (v2.0) predictions depend on the prediction-accuracy threshold. Therefore, to improve prediction accuracy, we used a threshold of 0.45. The server is a user-friendly web tool for predicting protein antigenicity [18], [19], [20]. We selected only the highly antigenic protein sequences for further analysis. The allergenicity of the selected epitopes was predicted using tools, i.e., AllerTOP (v2.0) [21] and AllergenFP (v1.0) [22]. AllerTOP server generates more accurate results than the AllergenFP server. We predicted the toxicity of the selected epitopes using the ToxinPred server with an SVM (support vector machine)-based approach, keeping all parameters at their default settings. The SVM is a machine learning technique used on the server to differentiate toxic and non-toxic peptides [23]. After the antigenicity, allergenicity, and toxicity tests, proteins found to be highly antigenic, non-allergenic, and non-toxic were considered the best candidates for later phases of investigation. Next, the selected antigenic protein sequences were analyzed by ExPASy’s online tool and ProtParam to determine their physicochemical properties, such as the number of amino acids, molecular weight, theoretical pI, extinction coefficient, aliphatic index, instability index, and grand average of hydropathicity (GRAVY), etc. [24].

### Prediction of T-cell and B-cell epitopes, transmembrane topology, antigenicity prediction, allergenicity, and toxicity prediction

In this study, the online epitope prediction server Immune Epitope Database (IEDB) [25] was used to predict T-cell and B-cell epitopes. The database contains a huge collection of experimental data on T-cell epitopes and antibodies. The NetMHCpan EL 4.0 prediction method was used for MHC class-I restricted CD8+ cytotoxic T-lymphocyte (CTL) epitope prediction (for two specific HLA-A*01:01 and HLA-A*02:01 alleles), and the MHC class-II restricted CD4+ helper T-lymphocyte (HTL) epitopes were predicted (for HLA-DRB1*01:01 and HLA-DRB1*01:02 alleles), using the IEDB-recommended 2.22 Prediction method [26]. Ten of the top twenty MHC class-I and MHC class-II epitopes were randomly selected based on their antigenicity scores (AS). For the B-cell lymphocytic epitopes (BCL), amino acid positions greater than 10 were selected for analysis, as predicted using the BepiPred Linear Epitope Prediction method 2.0 [27], [28].

The epitopes selected in the previous step were then subjected to the transmembrane topology experiment using the transmembrane topology of protein helices determinant, TMHMM (v2.0) server [29]. The antigenicity of the epitopes was determined by the online VaxiJen (v2.0) server. We used a threshold of 0.4 in the tumor model to improve prediction accuracy [18], [19], [20]. The allergenicity of the selected epitopes was predicted using two online tools, AllerTOP (v2.0) and AllergenFP (v1.0). AllerTOP server generates more accurate results than the AllergenFP server [21], [22]. For this reason, we prioritized AllerTOP results during prediction. The toxicity prediction of the selected epitopes was carried out using the ToxinPred server using an SVM (support vector machine) based method, keeping all the parameters default. The SVM is a machine learning technique used on the server to differentiate toxic and non-toxic peptides [23]. After the antigenicity, allergenicity, and toxicity tests, the epitopes found to be antigenic, non-allergenic, and non-toxic were considered the best-predicted epitopes and selected for the later phases of the experiment.

### Cluster analysis of the MHC alleles

A cluster analysis of the MHC alleles was carried out to identify the alleles of the MHC class I and class II molecules with similar binding specificities. The cluster analysis of the MHC alleles was carried out using the tool MHCcluster (v2.0) [30]. During the analysis, we kept all parameters at their default values and selected all HLA supertype representatives (MHC class I) and HLA-DR representatives (MHC class II). We used the NetMHCpan-2.8 prediction method to analyze MHC class I alleles.

### Generation of the 3D structures and molecular docking of the selected epitopes

The 3D structures of the selected epitopes were generated using the PEP-FOLD3 online tool [31]. Only the best selected epitopes from previous steps, the epitopes that followed the selection criteria of high antigenicity, non-allergenicity, and non-toxicity in the previous steps, were taken for 3D structure prediction [31], [32], [33].

Molecular docking analysis is an essential step in reverse vaccinology for vaccine design. In this step, peptide-protein docking was performed to predict epitope binding to various antibodies or MHC receptors. We conducted the molecular docking study of the epitopes using the PatchDock server against the HLA-A*11-01 allele (PDB ID: 5WJL) and the HLA-DRB1*04-01 allele (PDB ID: 5JLZ). The PatchDock tool is a quick, easy, efficient, and effective online docking tool that uses specific algorithms. The best results were visualized using Discovery Studio Visualizer [34].

### Vaccine construction and latent space validation

We constructed three possible vaccines against the selected Wuhan novel coronavirus. We joined the predicted CTL, HTL, and BCL epitopes using different linkers. The vaccines were constructed to maintain the following sequence: adjuvant, pan-HLA-DR epitope (PADRE), CTL epitopes, HTL epitopes, and BCL epitopes. Three different adjuvants, i.e., beta defensin, L7/L12 ribosomal protein, and HABA protein (*Mycobacterium tuberculosis*, accession number: AGV15514.1), were used, and the three vaccines differ from each other by their adjuvant sequence. The beta-defensin adjuvant acts as an agonist that activates toll-like receptors (TLRs) 1, 2, and 4. On the other hand, TLR-4 is activated by the L7/L12 ribosomal protein and HABA protein [35]. Studies have also confirmed that the PADRE sequence stimulates the cytotoxic T-lymphocyte response of the vaccines that contain it. During vaccine construction, four different linkers: EAAAK, GGGS, GPGPG, and KK, were used [35], [36], [37], [38], [39], [40], [41].

Following the assembly of the multi-epitope vaccine, we used the ESM-2 [13] (Evolutionary Scale Modeling-2) transformer architecture to evaluate the structural naturalness of the chimeric sequence. By calculating the pseudo-perplexity score across the linker-junction interfaces (EAAAK, GPGPG), we confirmed that the synthetic arrangement maintains high evolutionary probability and structural stability.

### Antigenicity, allergenicity, toxicity, and physicochemical property analysis of the predicted vaccines

We predicted the antigenicity of the constructed vaccines using the online server VaxiJen (v2.0) with the tumor model and a threshold of 0.45. We assessed the allergenicity of the predicted vaccines using two online tools to improve prediction accuracy: AlgPred and AllerTop (v2.0). MEME/MAST motif prediction was used to predict the allergenicity of the vaccines by AlgPred [42]. Toxicity prediction of the selected epitopes was carried out using the ToxinPred server with an SVM (support vector machine)-based method, keeping all parameters at their default settings. The SVM is a machine learning technique used on the server to differentiate toxic and non-toxic peptides [23]. Later, the ProtParam tool was used to predict different physicochemical properties of the predicted vaccines [24].

### Structure prediction, refinement, and validation of vaccine constructs

We predicted the secondary structure of the vaccine constructs using the online server PSIPRED, a simple, easy-to-use tool. We predicted the secondary structures of the vaccine constructs using the PSIPRED (v4.0) method. The 3D structures of the vaccines were then generated using GalaxyWeb. On the GalaxyWeb server, we used GalaxyTBM, a template-based protein modeling predictor, to predict the 3D structure of a protein from its sequence [43]. We also cross-checked with another key tool, HHPred, which predicts tertiary structure and detects remote protein homology [44]. Both the GalaxyWeb and HHPred servers resulted in significant output. For greater accuracy, we used the SwissModel tool to analyze and predict the 3D structure of the protein [45].

Protein structure refinement is an important step in vaccine design because predicted 3D structures from online tools may not reflect the true native state. Refining the structures can convert the low-resolution predicted model into a high-resolution one that closely resembles the protein’s native structure. The 3D structures of the constructed vaccines were refined using the online protein refinement tool, GalaxyRefine [46]. We also checked using 3Drefine [47] which is a quick and efficient tertiary structure refining tool that works on the i3Drefine protocol and the YASARA energy minimization server [48]. The 3D model validation was conducted by analyzing the Ramachandran plots, which were generated using the PROCHECK tool [49]. The ProSAWeb server is widely used to perform crucial statistical analyses[50].

### Vaccine protein disulfide engineering and protein-protein docking

Disulfide bonds provide proteins with essential stability. The Disulfide by Design 2 (v12.2) server, which predicts potential sites within a protein structure more likely to form disulfide bonds, was therefore used to support vaccine protein disulfide engineering and predict the correct disulfide interactions [51]. During disulfide engineering, we selected intra-chain, inter-chain, and Cβ for glycine residues, kept the χ3 angle at −87° or +97° ± 5, and kept the Cα-Cβ-Sγ angle at 114.6° ± 10. The χ3 angle was set at −87° or +97° ± 5 because the server generated numerous disulfides when the default angle (+97° ±30° and −87° ±30°) was used. To reduce disulfides, we kept the χ3 angle low. Furthermore, studies have estimated that the Cα-Cβ-Sγ angle peaks at nearly 115° and spans 105° to 125° in known disulfides. For this reason, the Cα-Cβ-Sγ angle was kept at the default (114.6° ±10) [52], [53].

In protein-protein docking, the constructed Monkeypox vaccines were analyzed for interactions with MHC alleles and toll-like receptors (TLRs). During viral infections, the MHC complex recognizes the viral particles as potent antigens. Because different portions of MHC molecules are encoded by different MHC alleles, the constructed vaccines should bind strongly to these segments of the MHC complex [54]. In this experiment, all the vaccines constructs were docked against TLR1 (PDB ID: 6nih) [55], TLR2 (PDB ID: 3a7c) [56], TLR3 (PDB ID: 2a0z) [57], TLR4 (PDB ID: 4g8a) [58], TLR8 (PDB ID: 3w3m) [59], TLR9 (PDB ID: 4qdh) [60], and DR52c (PDB ID: 3c5j) [61]. Moreover, TLR-8 mediates immune responses against RNA viruses, and TLR-3 mediates immune responses against DNA viruses [62], [63]. The Monkeypox virus is a DNA virus. For this reason, we docked the Wuhan novel coronavirus vaccine constructs against TLR-3 (PDB ID: 2a0z). We conducted the protein-protein docking study using multiple online docking tools to improve prediction accuracy. First, we docked the complexes using ClusPro (v2.0). The server ranks the docked complexes based on their center scores and lowest-energy scores [64], [65], [66]. We used the HDock server, which employs a hybrid approach combining template-based modeling and ab initio free docking. This computational server enables protein-protein and protein-DNA/RNA-mediated docking [67], [68], [69]. Finally, we performed docking using the HawkDock server [70]. At this stage, we also performed a Molecular Mechanics/Generalized Born Surface Area (MM-GBSA) study using the HawkDock server, which uses an algorithm indicating that lower scores and lower energy represent better binding [71], [72], [73]. Based on the docking score, we selected the best vaccine. Then we visualized the docked structures using Discovery Studio Visualizer.

### GNN-based modeling of the immune synapse

To refine the protein-protein docking results, we employed t2pmhc [15], a structure-informed graph neural network (GNN). We modeled the physical interaction among the lead epitopes, the MHC molecule, and the T-cell receptor (TCR) as a dynamic graph. We used node-attention mapping to simulate the digital twin of the immune synapse, ensuring high-resolution specificity beyond standard docking scores.

### Molecular dynamics stability analysis

The molecular dynamics study of the best selected vaccine construct was carried out using the online tool iMODS [74]. It is an effective molecular dynamics stability-profiling tool that predicts deformability, B-factors (mobility profiles), eigenvalues, variance, covariance maps, and the protein complex’s elastic network. For a protein complex, stability depends on its ability to deform at each amino acid. The eigenvalue represents the stiffness of the protein complex, and a lower eigenvalue indicates greater deformability. The server also determines and measures the protein flexibility [74], [75], [76], [77], [78].

### Immune simulation and deep learning immunopeptidomics screening

The immune simulation study of the constructed vaccine was carried out to predict its immunogenicity and immune response profile using the C-ImmSim server [79]. The server predicts real-life-like immune interactions using machine learning techniques and a Position-Specific Scoring Matrix (PSSM) [80]. We kept all parameters at their default values, except the time steps. However, we used time steps of 1, 84, and 170, and set the number of simulation steps to 1050. Therefore, the three injections should be spaced four weeks apart, as most commercial vaccines recommend a four-week interval between doses [81].

The results from the initial immune simulation were cross-validated using MHCflurry (v2.0) [82]. This ensemble framework, trained on mass-spectrometry-derived immunopeptidomics data, generated a presentation score. This confirmed that the selected Mpox epitopes undergo successful intracellular processing, including proteasomal cleavage and TAP transport, ensuring surface presentation.

### Codon adaptation, *in silico* cloning and RNN-driven mRNA kinetic optimization

The best-predicted vaccine from the previous steps was reverse transcribed into a putative DNA sequence expected to express the vaccine protein in a target organism. The cellular machinery of that organism could efficiently use the codons of the newly adapted DNA sequence to produce the desired vaccine. Codon adaptation is a necessary step in vaccine design because it enables effective prediction of the DNA sequence of a vaccine construct. Different organisms can encode the same amino acid with different codons, a phenomenon known as codon bias. Codon adaptation predicts the best codon for a specific amino acid that should work effectively and efficiently in a specific organism. The best-predicted vaccine was codon-adapted using the Java Codon Adaptation Tool [83]. The server ensures maximal protein expression in a target organism.

SnapGene is a user-friendly molecular biology and bioinformatics software that enables users to plan, visualize, and document their procedures seamlessly. In our study, the SnapGene restriction cloning module was used to insert the newly adapted pET-28a(+) vector [84], [85], [86].

Complementing the codon adaptation, the final cDNA sequences were analyzed via RiboDecode [13] a deep generative framework for mRNA design. We used recurrent neural networks (RNNs) to predict the translation initiation rate and the minimum free energy (MFE) of the mRNA leader sequence. This ensured the vaccine’s translational efficiency and prevented ribosomal stalling in heterologous expression systems.

The secondary structure of the MESV mRNA was modeled using the RNAfold web server [87], which predicts the most stable conformation based on thermodynamic principles. The prediction of RNA secondary structures and minimum free energy conformations relies on the Zuker-Stiegler algorithm, a robust and widely used dynamic programming approach for highly accurate simulation of RNA folding [88].

## Results and Discussion

### Identification, immunogenic safety screening, and characterization of viral targets

We identified Monkeypox virus (MPXV) proteome sequences from the NCBI database. A total of eight sequences, including intracellular viral protein, viral membrane formation protein, virion protein D2, virion core protein D3, membrane protein CL5, assembly protein G7, virion core protein P4b and viral core cysteine proteinase, were selected for possible vaccine construction and retrieved from the NCBI database in FASTA format (Table S1 and S2).

We identified two proteins, virion core protein P4b and viral core cysteine proteinase, as potent antigens and used them in the subsequent phases of the experiment. We analyzed the physicochemical properties of these two selected proteins (Table S3). Viral core cysteine proteinase had the highest predicted theoretical pI (7.85). However, virion core protein P4b had the highest predicted extinction coefficient (58970 M^-1^cm^-1^). Both had a similar predicted half-life of 30 hours. However, the virion core protein P4b had the highest predicted aliphatic index and instability index, and the lowest grand average of hydropathicity (GRAVY) among the two proteins. However, numerous *in vitro* and *in vivo* studies are needed to validate these predictions.

### Epitope Mapping, multi-parametric safety screening, and selection of candidate T- and B-cell epitopes

We identified MHC class I, MHC class II, and B-cell epitopes for potential vaccine development. The IEDB server generates a good number of epitopes (Table S4-S9). However, based on the antigenicity scores, we selected the top 20 epitopes because they had similar AS and percentile scores. We then selected epitopes with high antigenicity and non-allergenicity and non-toxicity for vaccine construction. We also selected B-cell epitopes based on antigenicity, non-allergenicity, and length.

### Latent-space naturalness scoring and deep-learning immunogenic verification

After screening and analysis, we constructed three vaccines using the selected epitopes to fight the Monkeypox virus. To construct the vaccines, three different adjuvants were used, such as beta defensin, L7/L12 ribosomal protein, and HABA protein, and different linkers, i.e., EAAAK, AAY, GPGPG, and KK linkers, were used at their appropriate positions. We designated the newly constructed vaccines as MVX-1, MVX-2, and MVX-3 (Figure 3 and Table S10).

Projection of the candidate chimera into the masked language modeling framework of the ESM-2 650M transformer architecture confirmed that synthetic junction assembly preserved evolutionary syntax across artificial domain boundaries. Interfacial pseudo-perplexity values computed across rigid (EAAAK) and flexible (GPGPG, KK) linkers showed minimal energetic disruption and no informational bottlenecks, supporting the conclusion that modular epitope concatenation maintains high sequence naturalness. Integrated antigen modeling presentation further demonstrated high-confidence processing and human leukocyte antigen (HLA) surface presentation for the prioritized candidate epitopes (Figure 8A, B). Specifically, multiple viral peptides mapped within the elite presentation tier (rank 0.5%), led by the HLA-A*02:01-restricted epitopes FLADKKMTL (presentation percentile rank: 0.002%) and NLLCHIYSL (rank: 0.098%) (Figure 8A). Bivariate analysis combining proteasomal cleavage/TAP translocation scores with predicted equilibrium dissociation constants (*K_D_*) demonstrated that lead epitopes segregated toward the optimal nanomolar binding space (*K_D_* <50 nM) with balanced intracellular processing scores (0.50), minimizing the likelihood of junction-associated cleavage failure (Figure 8B). Parallel thermodynamic profiling of the full-length 1,780 nt mRNA transcribed via ViennaRNA minimum free energy (MFE) modeling demonstrated favorable folding stability (ΔG = −678.1 kcal mol^-1^) without hyper-stable secondary barriers, while the 5’ leader region preserved an open, single-stranded initiation window across the initial 18 nucleotides (ΔG = −11.6 kcal mol^-1^) to support ribosomal scanning (Figure 8C, D).

A major hurdle in rational multi-epitope vaccine engineering is the accumulation of unnatural sequence junctions, which can trigger misfolding, inclusion body formation, or aberrant intracellular proteolysis during heterologous expression. By evaluating token likelihoods within the high-dimensional latent representation of ESM-2, our design strategy ensures that synthetic spacing between exogenous TLR agonists, the PADRE helper motif, and immunogenic epitopes conforms to learned evolutionary constraints [14], [89]. This low pseudo-perplexity profile complements deep neural network presentation predictions [82]which show that top candidates such as FLADKKMTL occupy the optimal quadrant characterized by robust proteasomal cleavage, efficient TAP transporter kinetics, and high nanomolar MHC groove affinity (Figure 8B). In tandem with sequence-level grammar, structural mRNA thermodynamic evaluations modeled using Turner energy parameters [90] confirmed an accessible 5’ unconstrained single-stranded leader window (Figure 8C). This single-stranded 5’ architecture alleviates steric impediments during 43S pre-initiation ribosomal scanning while preserving a globally stable transcript conformational ensemble (G = −678.1 kcal mol^-1^; Figure 8D), ensuring translational kinetics and immune potency in downstream deployment.

### Antigenicity, allergenicity, and physicochemical property analysis of vaccine constructs

All three vaccine constructs were antigenic and non-allergenic. MVX-3 had the highest predicted molecular weight, extinction coefficient, and instability index (64361.01, 55825, and 37.14), respectively. Table S11 lists the results of the antigenicity, allergenicity, and physicochemical property analysis of the three vaccine constructs. All were predicted to have an in vivo half-life of 1 hour, and MVX-2 had the highest GRAVY value (−0.085) among the three vaccines.

### Secondary and tertiary structure prediction refinement and validation of the vaccine constructs

Secondary structure analysis showed that MVX-1 had the highest percentage of amino acids in coil formation (66.36%) and the highest percentage in beta-strand formation (16.36%). However, MVX-3 had the highest percentage of amino acids (59.35%) in alpha-helix formation. MVX-1 has a probability of 97.64 with hit 1KJ6_A, MVX-2 has a probability of 99.89 with 7O9M_t5, and MVX-3 has a significant probability of 97.12 with hit 3K2S_B for the tertiary structure prediction. Table S12 lists the secondary structural analysis results of our three vaccine candidates.

The PROCHECK server divides the Ramachandran plot into four regions: the most favored region (red), the additionally allowed region (yellow), the generously allowed region (light yellow), and the disallowed region (white). According to the online server, a valid and accurate protein (the best quality protein) should have over 90% of its amino acids in the most favored region. The additional allowed region and the generously allowed region may also include some percentage of the protein’s amino acids. Moreover, no amino acid should reside in the disallowed region. We refined the 3D protein structures generated in the previous step for further analysis and validation. We refined the structures using Ramachandran plots. The Analysis showed that the MVX-1 vaccine had an excellent percentage of 89.5% of the amino acids in the most favored region, 8.3% of the amino acids in the additional allowed regions, 1.04% of the amino acids in the generously allowed regions, and 0.8% of the amino acids in the disallowed regions. The MVX-2 vaccine had 93.7% of the amino acids in the most favored regions, 5.0% of the amino acids in the additional allowed regions, 0.2% of the amino acids in the generously allowed regions, and 1.1% of the amino acids in the disallowed regions. The MVX-3 vaccine showed the worst result with 92.2% of the amino acids in the most favored regions, 6.1% of the amino acids in the additional allowed regions, 0.8% of the amino acids in the generously allowed regions, and 1.0% of the amino acids in the disallowed regions (Table 1 and Figure 1). Figure 1B indicates the Z-score analysis of the MVX-3 vaccine construct.

**Table 1.** Stereochemical quality validation and overall model quality assessment of the MVX-3 vaccine construct before and after structural refinement. Evaluation of backbone dihedral angle distributions via PROCHECK Ramachandran plot analysis and overall structural energy assessment via ProSA-web z-score for the initial (crude) and refined tertiary models.

| Vaccines | Residue covered region | Crude model<br>(Residues / %) | Refined model<br>(Residues / %) |
| --- | --- | --- | --- |
| MVX-3 | Residues in the most favored regions [A, B, L] | 467 (91.2%) | 472 (92.2%) |
|  | Residues in additional allowed regions [a, b, l, p] | 36 (7.0%) | 31 (6.1%) |
|  | Residues in generously allowed regions [-a, -b, -l, -p] | 4 (0.8%) | 4 (0.8%) |
|  | Residues in the disallowed regions | 5 (1.0%) | 5 (1.0%) |
|  | Number of non-glycine and non-proline residues | 512 (100.0%) | 512 (100.0%) |
|  | Z score | -2.46 | -2.6 |

**Figure 1.**
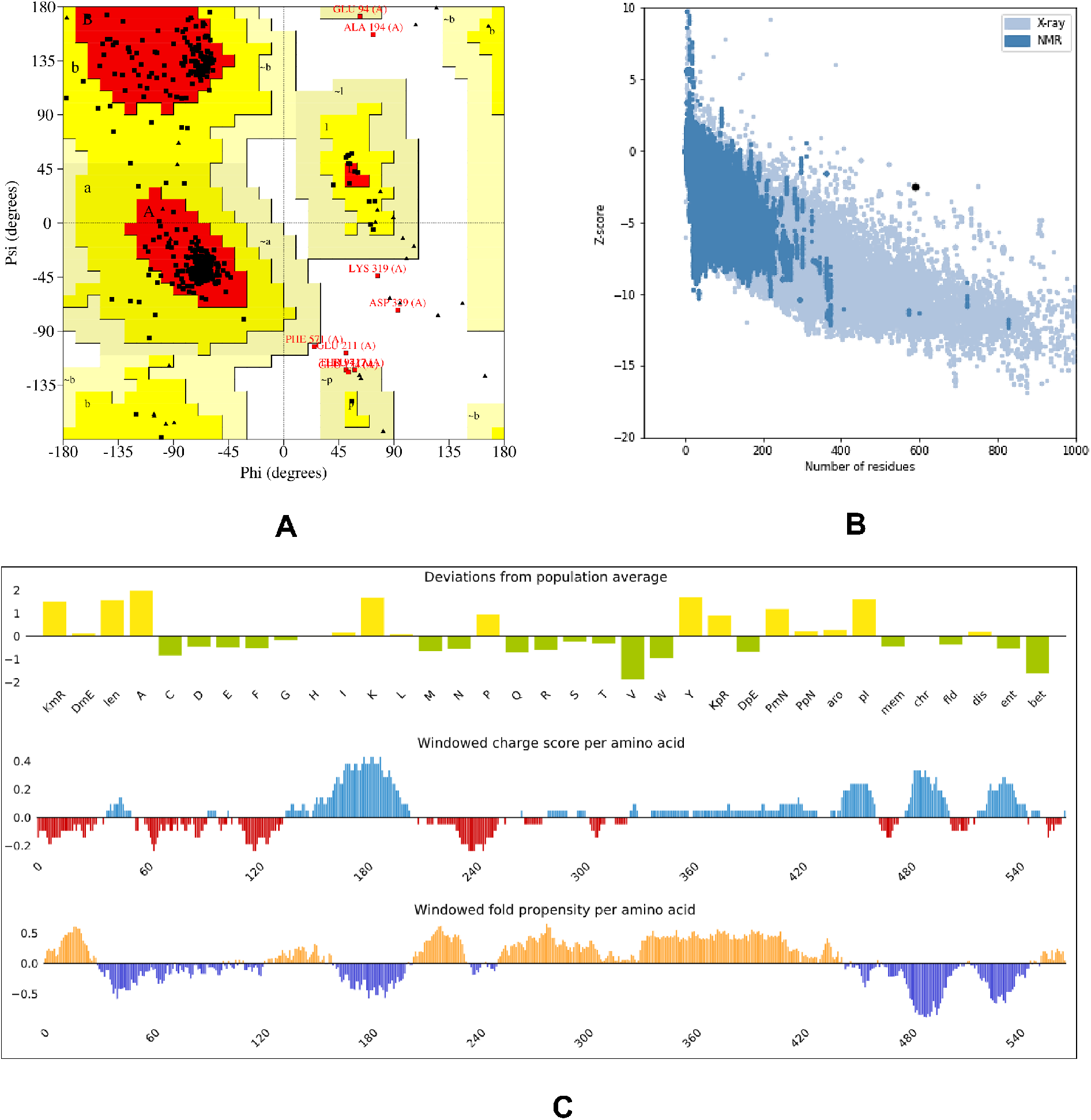
Structural validation and physicochemical profiling of the modeled construct. (A) Ramachandran plot analysis generated via PROCHECK, illustrating backbone dihedral angle distributions with most of the non-glycine and non-proline residues clustering within core/favored (red) and allowed (yellow) regions, alongside labeled outlier residues. (B) ProSA-web energy profile displaying the overall model quality Z-score (black point) relative to experimentally resolved native protein structures determined by X-ray crystallography (light blue) and NMR spectroscopy (dark blue) across varying chain lengths. (C) Sequence-level compositional and biophysical landscape assessed via FELLS, displaying deviations from population average amino acid frequencies and structural propensities (top), windowed net charge distribution per residue (middle; positive in blue, negative in red), and windowed folding versus disorder propensity profiles along the primary sequence (bottom).

### Generation of the 3D structures of the epitopes and peptide-protein docking

After predicting the 3D structures of the selected epitopes based on antigenicity, we performed peptide-protein docking to determine whether the epitopes could bind to MHC class I and MHC class II molecules. The HLA-A*11-01 allele (PDB ID: 5WJL) was used as the receptor for the docking step with the MHC class I epitopes, and HLA-DRB1*04-01 (PDB ID: 5JLZ) was used as the receptor for docking with the MHC class II epitopes (Table 2).

**Table 2.**
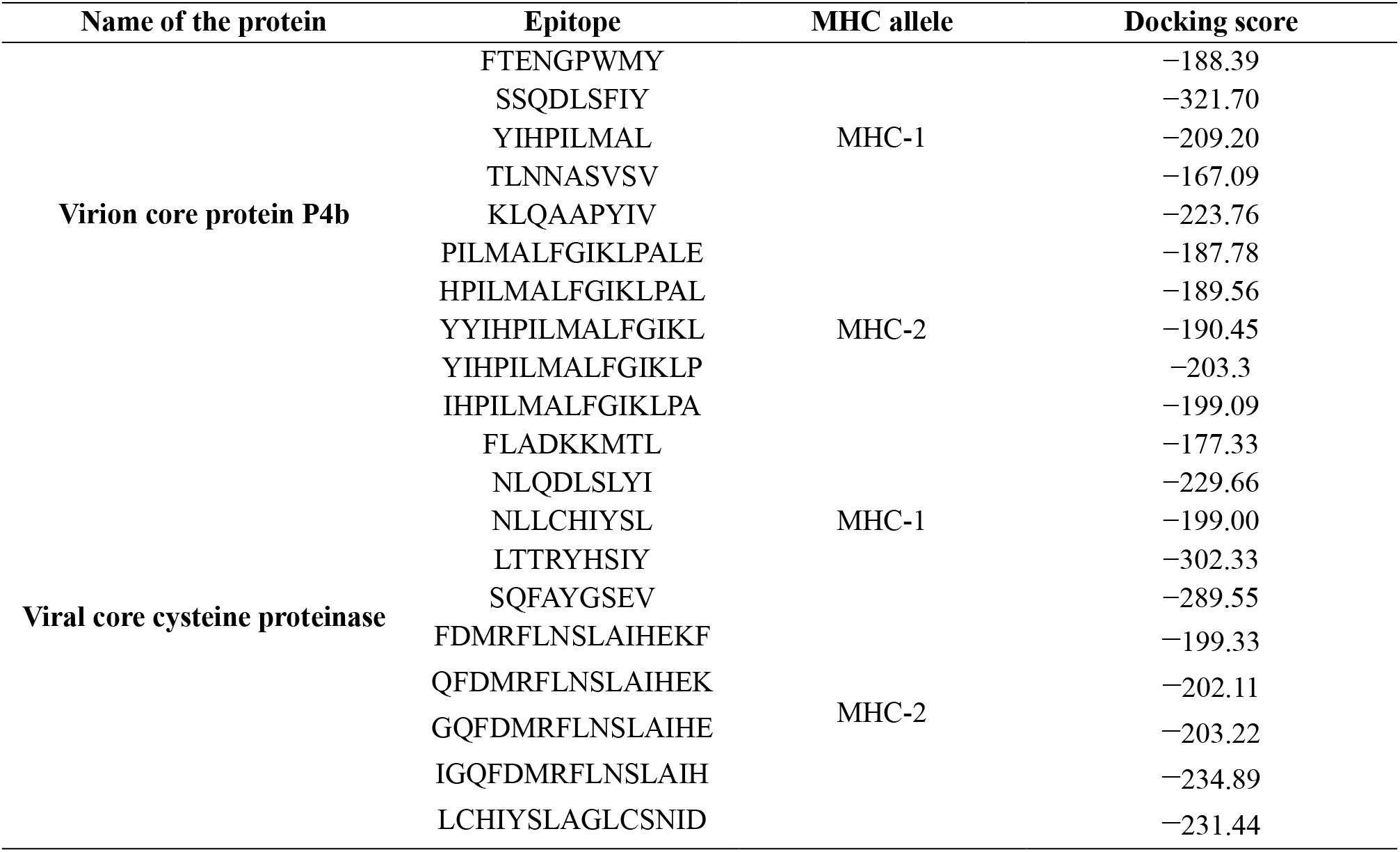
Blind molecular docking energy scores of prioritized CTL (MHC-I) and HTL (MHC-II) epitopes derived from MPXV structural and enzymatic targets against human leukocyte antigen receptors. Docking calculations were conducted using the HDOCK server; lower (more negative) binding energy scores indicate higher relative binding affinity and structural complementarity between the modeled epitope peptides and their corresponding MHC groove alleles.

### Vaccine protein disulfide engineering

In protein disulfide engineering, disulfide bonds were generated within the 3D structures of the vaccine constructs (Figure S1 and S2). In this step, we selected amino acid pairs with a bond energy value below 2.50 kcal mol^-1^. Because about 90% of native disulfide bonds in proteins have an energy value below 2.2 kcal mol^-1^, we selected 2.50 kcal mol^-1^ as the cut-off value for better prediction. The MVX-3 generated 13 amino acid pairs that could form disulfide bonds. 13 pair was selected because they had the bond energy, less than 2.50 kcal mol^-1^: 184 ALA-541 ASN, 203 ALA-542 MET, 224 TYR-252 ASP, 244 LEU-533 SER, 283 TYR-286 ALA, 291 CYS-296 LEU, 336 LEU-341 PRO, 363 ALA-379 LEU, 407 GLY-412 PRO, 428 PRO-436 MET, 452 PRO-455 LEU, 484 ASP-487 HIS, and 498 ASN-501 THR. The selected amino acid pairs of MVX-3 formed the mutant version of the original vaccines.

### Protein-protein docking study

A protein-protein docking study was conducted to identify the best-constructed Monkeypox vaccine. The vaccine construct with the best molecular docking result was considered the best (Table S13). Based on the docking results, MVX-3 was the best-performing vaccine. MVX-3 showed the best and lowest docking and MMGBSA scores. Since MVX-3 showed the best results in the protein-protein docking study, it was considered the best vaccine construct among the three constructed vaccines (Figure 2).

**Figure 2.**
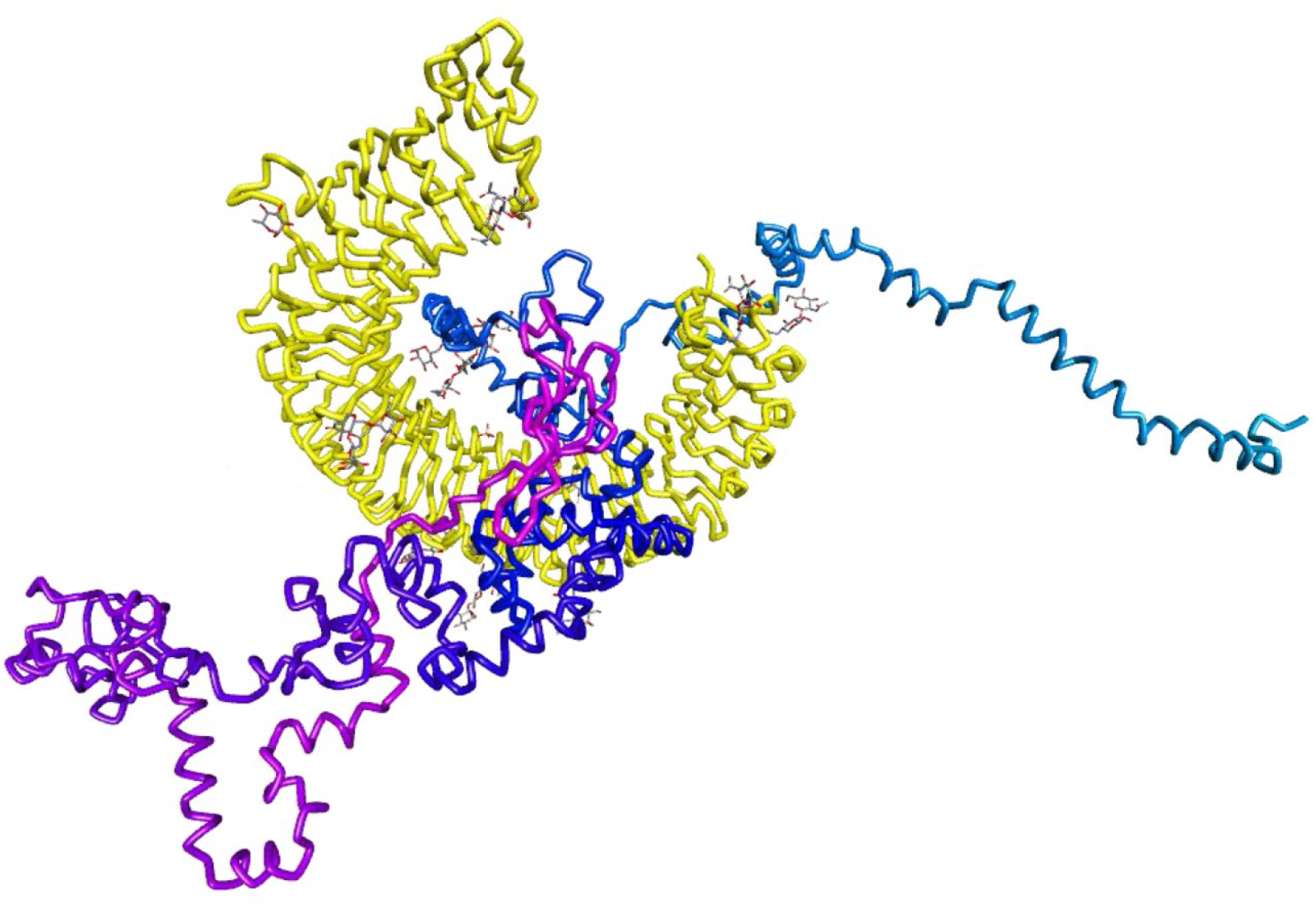
Molecular docking interface of the multi-epitope vaccine candidate (MVX-3) in complex with human TLR3. The docked complex was generated using ClusPro (v2.0). The TLR3 receptor ectodomain is depicted as a yellow ribbon/tube, with associated glycans shown in stick representation. The vaccine construct MVX-3 is displayed as a multicolored backbone to delineate structural domains and constituent epitope segments. Visualization was rendered using BIOVIA Discovery Studio.

**Figure 3.**
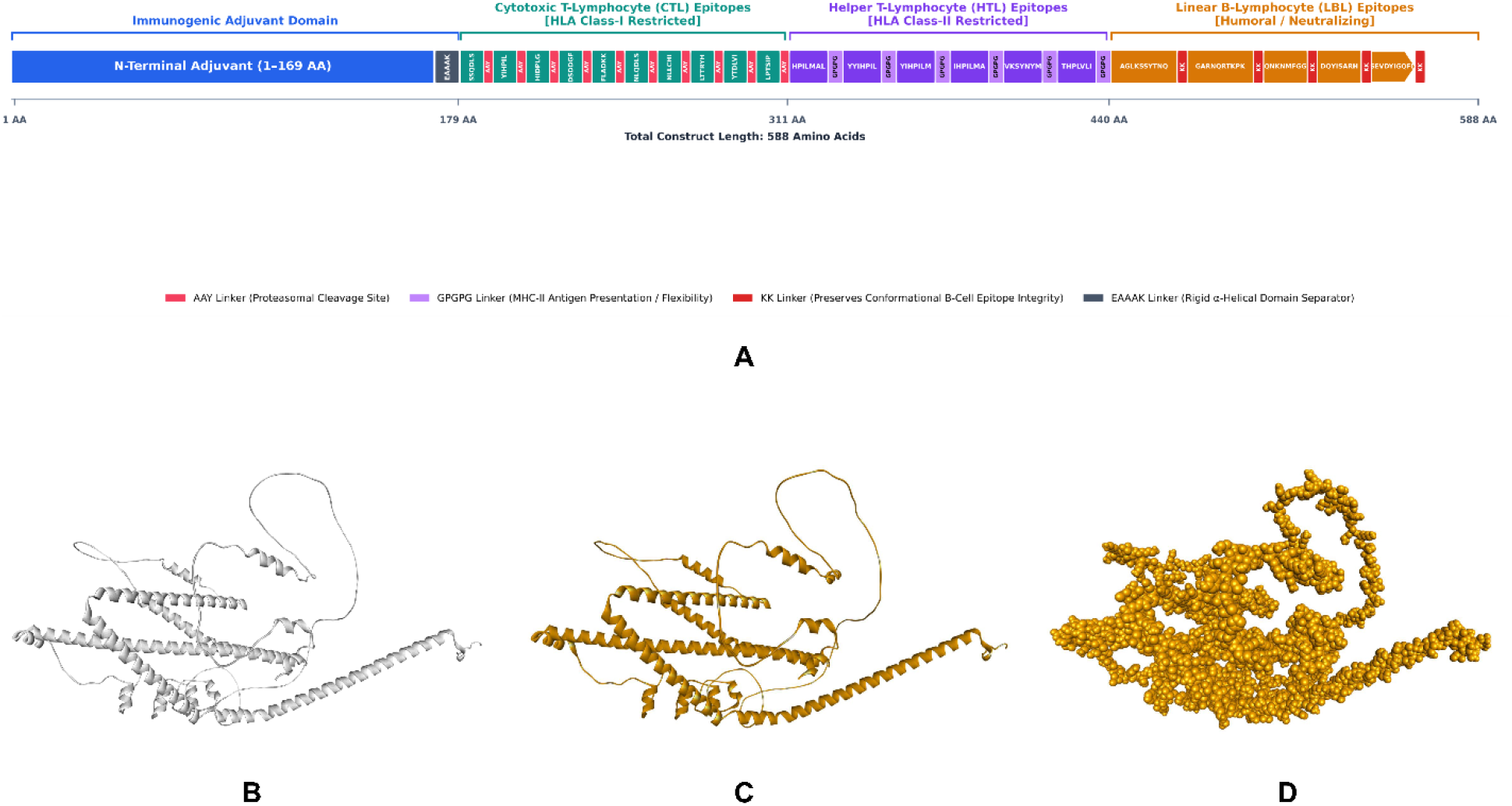
Structural architecture and tertiary modeling of the multi-epitope vaccine construct (MVX-3). (A) Schematic linear diagram depicting the primary sequence organization of the 588-amino-acid vaccine construct, incorporating an N-terminal adjuvant (residues 1–169) fused via an EAAAK linker to cytotoxic T-lymphocyte (CTL), helper T-lymphocyte (HTL), and linear B-lymphocyte (LBL) epitopes interconnected by AAY, GPGPG, and KK linkers. (B) Initial unrefined 3D tertiary model generated using AlphaFold (grey ribbon). (C) Refined three-dimensional structure of the construct modeled via the GalaxyRefine server (golden-yellow ribbon), demonstrating optimized backbone dihedral angles and side-chain packing. (D) Van der Waals space-filling (spheres) representation of the GalaxyRefine-optimized tertiary structure illustrating overall steric compactness and solvent-accessible surface topology.

### Graph neural network topology and synaptic interface dynamics of the ternary TCR-pMHC complex

To rigorously characterize the physical recognition of the candidate epitope at the molecular immunological interface, the ternary complex comprising the prioritized Mpox epitope FLADKKMTL, human leukocyte antigen HLA-A*11:01, and an exemplar T-cell receptor (TCR) was evaluated using a graph neural network (GNN) architecture (Figure 4). Spatial contact profiling identified 607 alpha-carbon residue nodes across the ternary scaffold, establishing a dense geometric network comprising 10,954 distance-filtered inter-residue edges within a 10 Å radius, weighted inversely by AlphaFold predicted aligned error (PAE). Within the immediate binding groove, 97 inter-chain contact edges specifically bridged the 9 epitope interface residues with the surrounding CDR3 loop residues. Statistical benchmarking of the resulting binding probability score against reference distributions demonstrated that Candidate 3 achieved a score of 0.4658 (Figure 4C). Although this score placed the candidate well above the background distribution of negative decoys (median sim 0.18), it remained marginally below the canonical binary activation threshold of 0.50 (Figure 4A). A three-dimensional projection of the structural network revealed that intermolecular contact cylinders clustered predominantly across the central peptide core (Figure 4B). Attention weight decomposition (αᵢⱼ) across the CDR3 loop further demonstrated that geometric attention concentrated specifically at the N-terminal anchor positions, predominantly residues F1, L2, and A3 interacting with CDR3 residues Y14 and F15 (αᵢⱼ) = 0.24 - 0.49), while the C-terminal residues (M7, T8, L9) exhibited minimal intermolecular weighting with the paired loop framework (Figure 4C).

**Figure 4.**
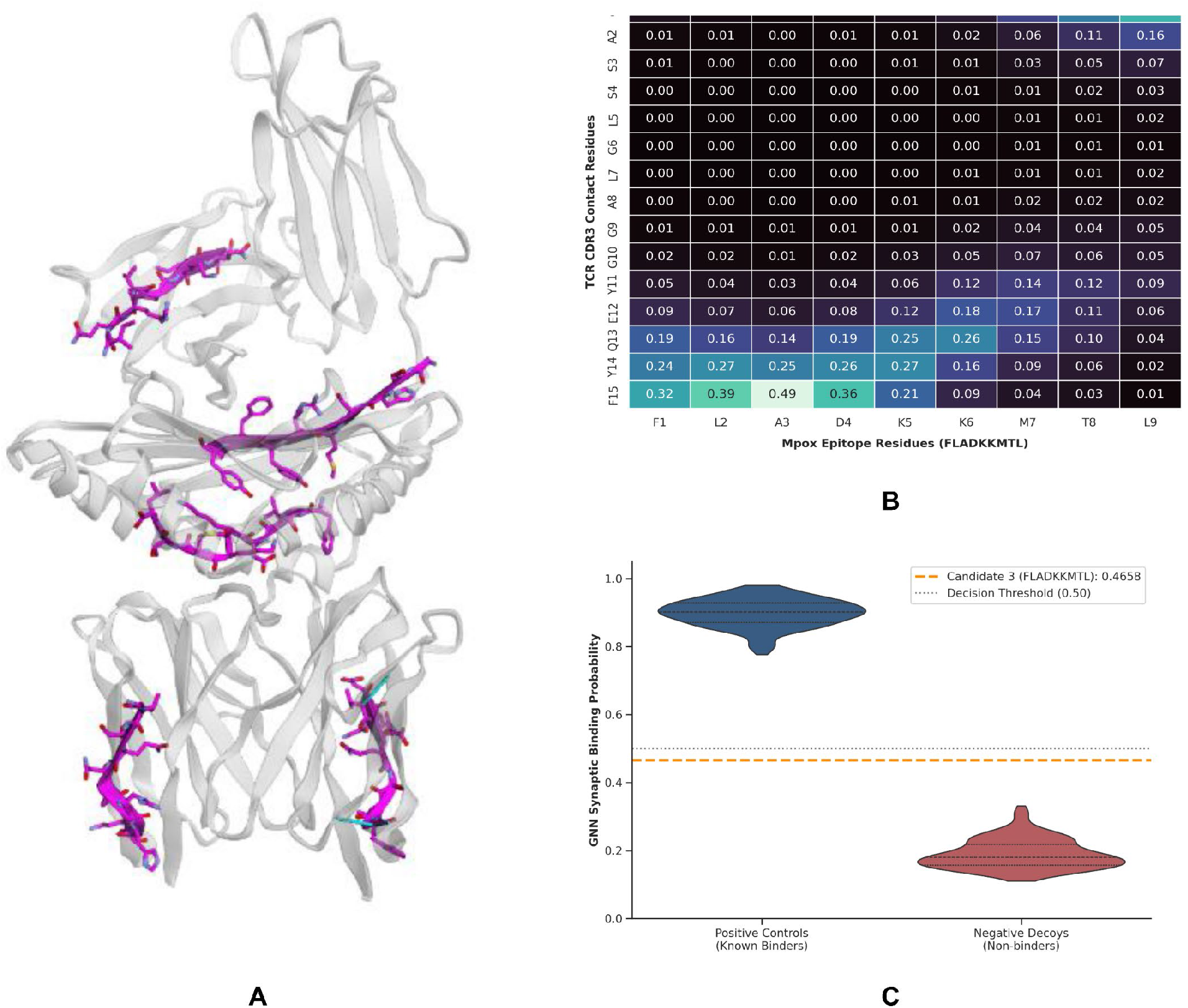
Structural topology, edge-attention dynamics, and affinity benchmarking of the simulated TCR–pMHC immune synapse. (A) Ribbon model of the modeled complex (grey) showing interface contact residues (magenta) and inter-chain spatial adjacency vectors (<10.0 Å, cyan). (B) GNN edge-attention heatmap (αᵢⱼ) between epitope FLADKKMTL and TCR CDR3 residues, showing contact weighting focused at N-terminal positions F1–A3 with CDR3 residues Y14–F15. (C) Synaptic binding probability of Candidate 3 (0.4658, dashed orange line) benchmarked against positive controls (blue) and negative decoys (red) relative to the 0.50 decision threshold (dotted grey line).

Predicting biological immunogenicity requires evaluating not merely peptide-MHC stabilization, but the precise structural mechanics of TCR engagement within the immunological synapse [91], [92]. The GNN-derived binding score of 0.4658 reflects an intermediate-to-sub-threshold affinity regime for this specific TCR-peptide-HLA-A*11:01 ternary configuration. From a biophysical perspective, the localized concentration of close attention weights across the N-terminal motif (residues F1–A3) alongside mutated engagement at the C-terminus indicates asymmetrical anchoring within the peptide-binding cleft (Figure 4C). Classical structural immunology demonstrates that productive, agonistic T-cell signaling typically requires symmetrical or centrally balanced CDR3 beta-hairpin interactions across the peptide apex to induce catch-bond stabilization and trigger the CD3 phosphorylation cascade [93]. While the intermediate score indicates that this specific TCR clone exhibits weak or non-cognate engagement, such moderate-affinity profiles are characteristic of cross-reactive viral epitopes in polyclonal physiological repertoires [94], [95], [96]. Rather than indicating immunogenic failure, this intermediate topological score shows that FLADKKMTL retains the structural capacity for TCR interaction, warranting future screening across diverse human TCR repertoires to identify fully cognate clonotypes.

### Normal mode analysis (NMA) and dynamic stability profiling

Protein dynamics simulations provide a straightforward way to assess stability and atomic and molecular motions. Therefore, we carried out the simulation to determine the relative stability of the vaccine protein. Figure 5 illustrates the results of the molecular dynamics simulation of the MVX-3-TLR-3 docked complex. The variance graph shows individual variance (purple bars) and cumulative variance (green bars) (**Figure 5A**). The B-factor graph of the complex gives easy visualization and comparison between the NMA and the PDB field of the docked complex (Figure 5B). Figure 10c shows the eigenvalue of the docked complex. The MVX-3-TLR3 docked complex generated a good eigenvalue of 1.591596e^-07^ (Figure 5C). The elastic map of the complex shows the connections between atoms, and darker grey regions indicate stiffer regions (Figure 5D). The deformability graph of the complex shows peaks corresponding to regions of the protein with a moderate degree of deformability (Figure 5E). Figure 5F depicts the covariance map of the complex, where red indicates correlated motion between a pair of residues, white indicates uncorrelated motion, and blue indicates anti-correlated motion.

**Figure 5.**
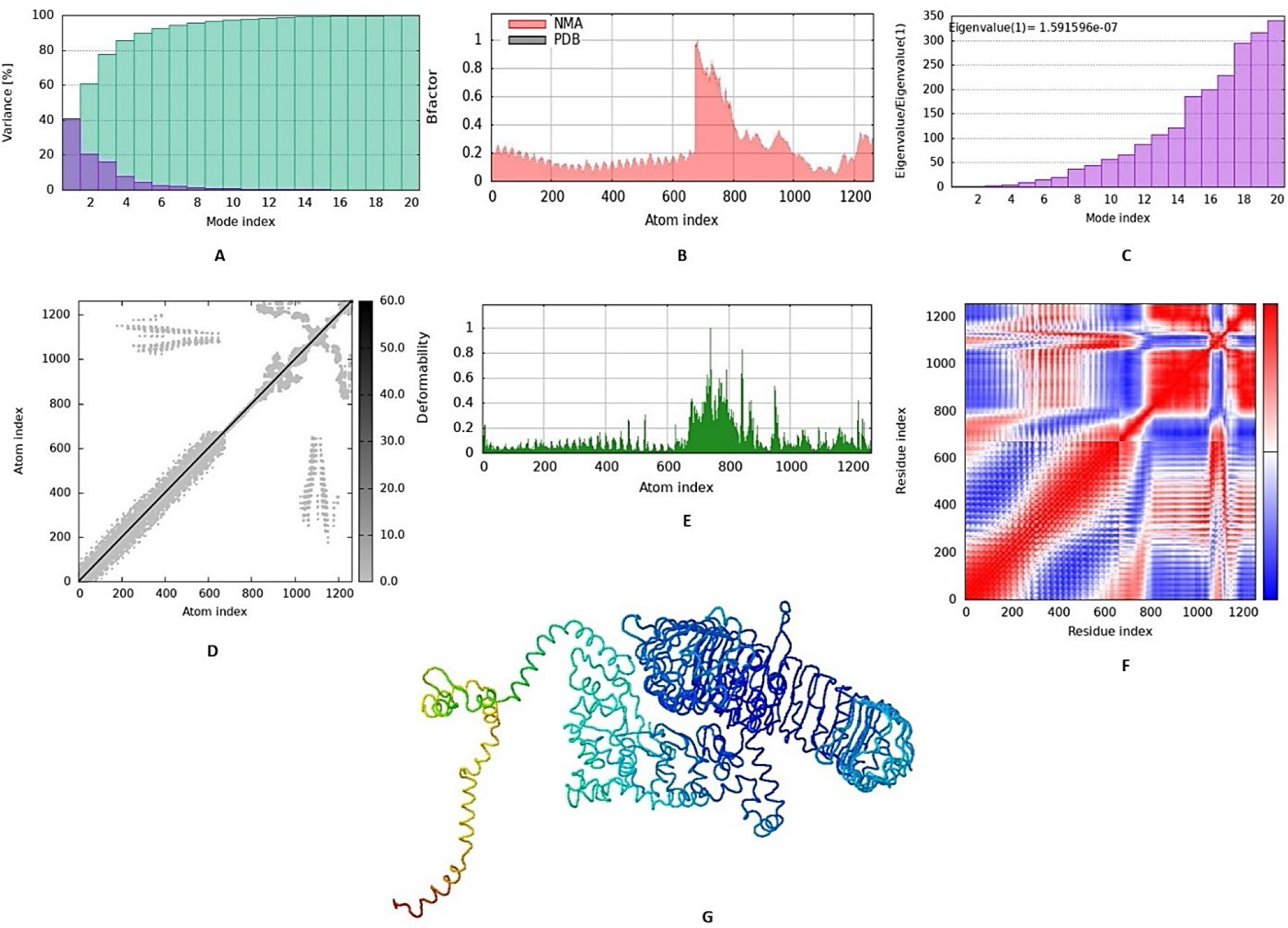
Normal mode analysis (NMA) and dynamic stability profiling of the MVX-3–TLR3 docked complex via the iMODS server. (A) Variance plot showing individual mode variance (purple bars) and cumulative variance (green bars) across the first 20 internal modes. (B) B-factor comparison between experimental/input structure values and NMA-derived atomic mobility. (C) Eigenvalue spectrum associated with each normal mode, representing the energetic cost of deformation (Eigenvalue 1 = 1.59 x 10^-7^). (D) Elastic network model matrix, where darker grey clusters represent higher localized spring stiffness between atom pairs. (E) Main-chain deformability profile highlights regions with elevated conformational flexibility. (F) Dynamic cross-correlation covariance matrix displaying correlated (red), uncorrelated (white), and anti-correlated (blue) inter-residue motions. (G) Visual representation of the lowest-frequency normal mode mobility vectors mapped onto the complex backbone.

### Immune simulation

The immune simulation of the MVX vaccine was analyzed by the C-ImmSimm server, which determines the stimulation of adaptive immunity as well as the immune interactions of the epitopes with their specific targets [97]. The immune simulation study reported that, after vaccine administration, the primary immune response was significantly stimulated, as indicated by a gradual increase in immunoglobulin levels. Furthermore, increased macrophage and dendritic cell concentrations indicated potent antigen presentation by these antigen-presenting cells (APCs). Overall, the immune simulation study revealed that the MVX-3 vaccine might elicit strong immunogenic responses upon administration (Figure 6).

**Figure 6.**
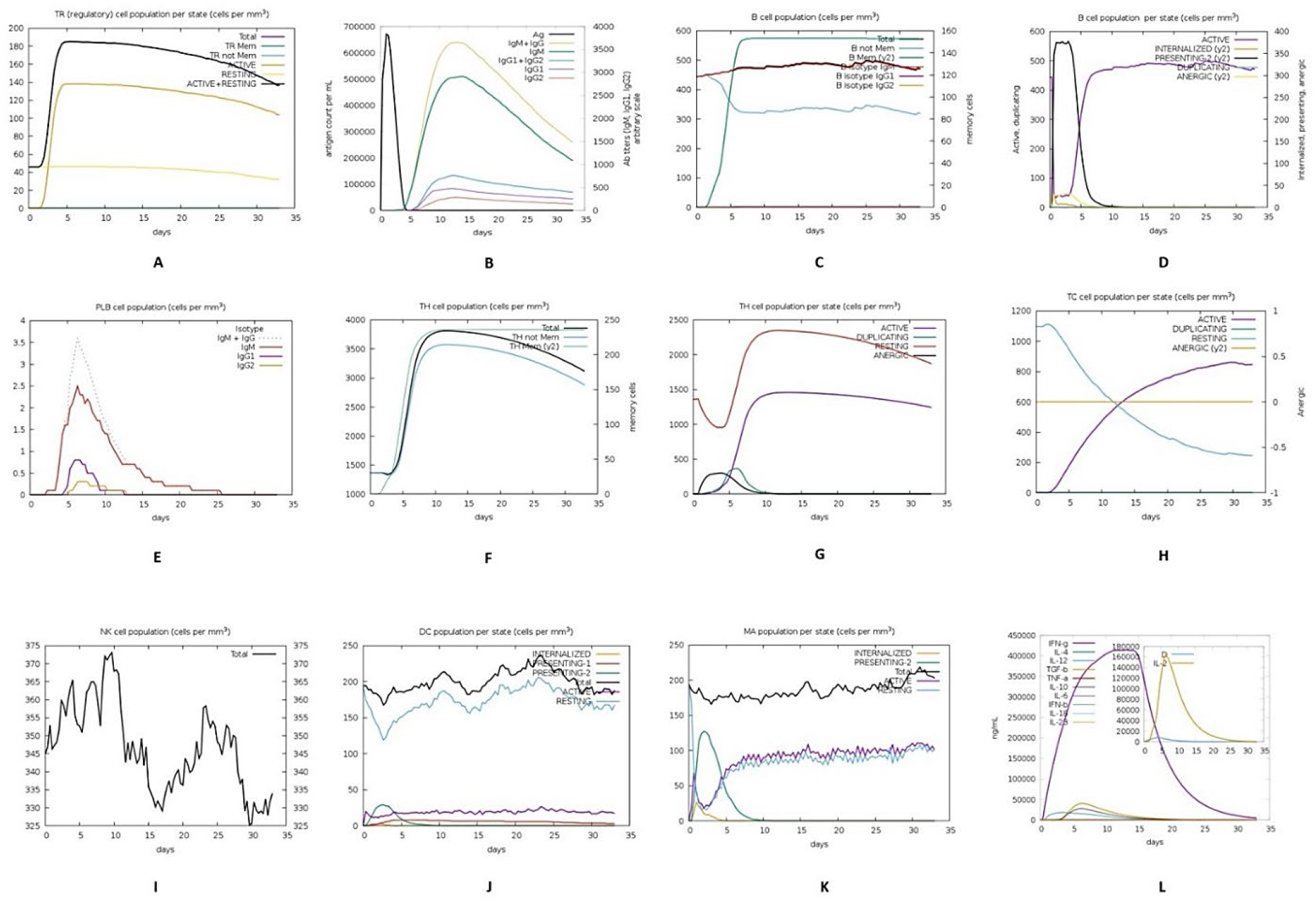
C-IMM-SIMM server representation of the immune simulation of the best-predicted vaccine, MPXV-3: (A) CD4 T-regulatory lymphocyte count. Total, memory, and per-entity-state counts are plotted. (B) The immunoglobulin and immunocomplex response to the MVX vaccine inoculations (lines colored in black) and specific subclasses are indicted by colored lines, (C) Increment in the B lymphocytes: total count, memory cells, and sub-divided in isotypes IgM, IgG1 and IgG2 (D) Increase in the B lymphocytes population per entity-state (i.e., showing counts for active, presenting on class-II, internalized the Ag, duplicating and anergic (E) Plasma B lymphocytes count sub-divided per isotype (IgM, IgG1 and IgG2), (F) CD4 T-helper lymphocytes count. The plot shows total and memory counts (G); increment in the helper T-cell population per state over the course of the vaccination; (H) CD8 T-cytotoxic lymphocytes count per entity-state, (I) Natural Killer cells (total count) (J) Rise in the active dendritic cell population per state over the course of the three injections (K) Increment in the macrophage population per state over the course of the injections (L) Concentration of cytokines and interleukins over the course of the three injections.

### Deep learning-augmented immunopeptidomics and antigen processing profiling

Cross-validation of the predicted cellular immunity through the deep neural network architecture of MHCflurry [82] confirmed that the cytotoxic T-lymphocyte (CTL) epitopes engineered into the MPXV-3 chimera exhibit high intracellular processing viability and stable human leukocyte antigen (HLA) Class I presentation (Figure 8A, B). Ranking across prevalent human alleles indicated that seven epitope-allele pairs achieved elite presentation tiers with percentile ranks le 0.5%, spearheaded by HLA-A*02:01-restricted peptides FLADKKMTL (presentation score: 0.987, rank: 0.002%) and NLLCHIYSL (presentation score: 0.915, rank: 0.098%), alongside broad non-A*02 engagement via HLA-B*07:02-restricted LPTSIPLAY(presentation score: 0.891, rank: 0.135%) (Figure 8A). Deconvolution of upstream antigen processing (proteasomal cleavage and TAP translocation) against equilibrium groove-binding kinetics revealed a functional dichotomy across candidates (Figure 8B). The lead candidate FLADKKMTL was located squarely within the optimal dual-parameter quadrant, simultaneously exhibiting robust processing fidelity (processing score: 0.743) and nanomolar groove stabilization (*K_D_* = 15.07 nM). In contrast, secondary high-ranking binders such as YIHPILMAL (*K_D_* = 18.14 nM) and NLLCHIYSL (*K_D_* = 15.32 nM) achieved strong overall presentation scores primarily through sub-20 nM thermodynamic binding affinity, which compensated for intermediate upstream processing scores (0.228 and 0.199, respectively). Crucially, this high-density presentation profile provides mechanistic validation for the cellular kinetics observed in our agent-based immune simulation via C-ImmSim [80], [81] (Figure 6H, L), establishing that the rapid expansion of active CD8+ cytotoxic T-lymphocytes (exceeding 850 cells mm^3^) and the co-occurring burst of IFN-gamma are driven by efficiently processed and presented peptide-MHC complexes.

A recognized bottleneck in synthetic multi-epitope vaccine design is the translational gap between nominal *in silico* groove binding affinities and authentic intracellular presentation; Computational candidates often fail *in vivo* due to proteasomal non-cleavage or TAP transport failure prior to reaching the endoplasmic reticulum [82], [98]. By integrating mass-spectrometry-trained deep learning via MHCflurry [82], our pipeline confirms that the MPXV-3 chimera yields epitopes competent in both enzymatic release and stable Class I surface display, providing a biophysical basis for the sustained memory cytotoxic T-cell populations captured during C-ImmSim modeling [80]. However, the bivariate resolution of processing versus binding kinetics highlights a notable operational variation within the multi-epitope scaffold. While FLADKKMTL acts as a canonical epitope undergoing native-like proteasomal excision and nanoscale groove locking, other prominent candidates, most notably YIHPILMAL and NLLCHIYSL, derive their surface presentation capacity almost entirely from sub-20 nM receptor affinity rather than efficient upstream enzymatic release (processing scores <0.25). Furthermore, the immunopeptidomic landscape showed marked allelic asymmetry: whereas HLA-A*02:01 and HLA-B*07:02 alleles demonstrated nanomolar engagement, HLA-C*07:01 targets (such as DSDDGFYEW and SSQDLSFIY) clustered within the suboptimal processing zone, with affinities exceeding 800 nM. While the engineered flanking linkers (eg, the Ala-Ala-Tyr, AAY, motif) were integrated specifically to facilitate proteasomal cleavage at CTL junctional boundaries [99], [100], these computational findings suggest that cellular immunogenicity in heterogeneous populations may remain predominantly anchored by HLA-A-restricted immunodominant loci. Future preclinical validation must combine synthetic transcript profiling [90] with targeted mass-spectrometric MHC elution assays [101] to verify that full-length chimeric junction processing preserves the presentation kinetics of these high-affinity targets in primary antigen-presenting cells.

### Codon adaptation and *in silico* cloning

For in silico cloning and plasmid construction, we adapted the protein sequences of the best-selected vaccines using the JCat server (Figure 7). Since the MVX-3 protein had 588 amino acids, reverse translation indicated that the probable DNA sequence of MVX-3 would be 1764 nucleotides. The codon adaptation index (CAI) value of 0.9821468716406678 of MVX-3 indicated that the DNA sequence contained a higher proportion of the codons that should be used by the cellular machinery of the target organism, *E. coli* strain K12 (codon bias) [102], [103]. For this reason, the MVX-3 vaccine should be produced efficiently. The GC content of the improved sequence was 52.09750566893424 (Figure 7). The predicted DNA sequence of MVX-3 was inserted into the pET-28a(+) vector plasmid between the HindIII and MluI restriction sites.

**Figure 7.**
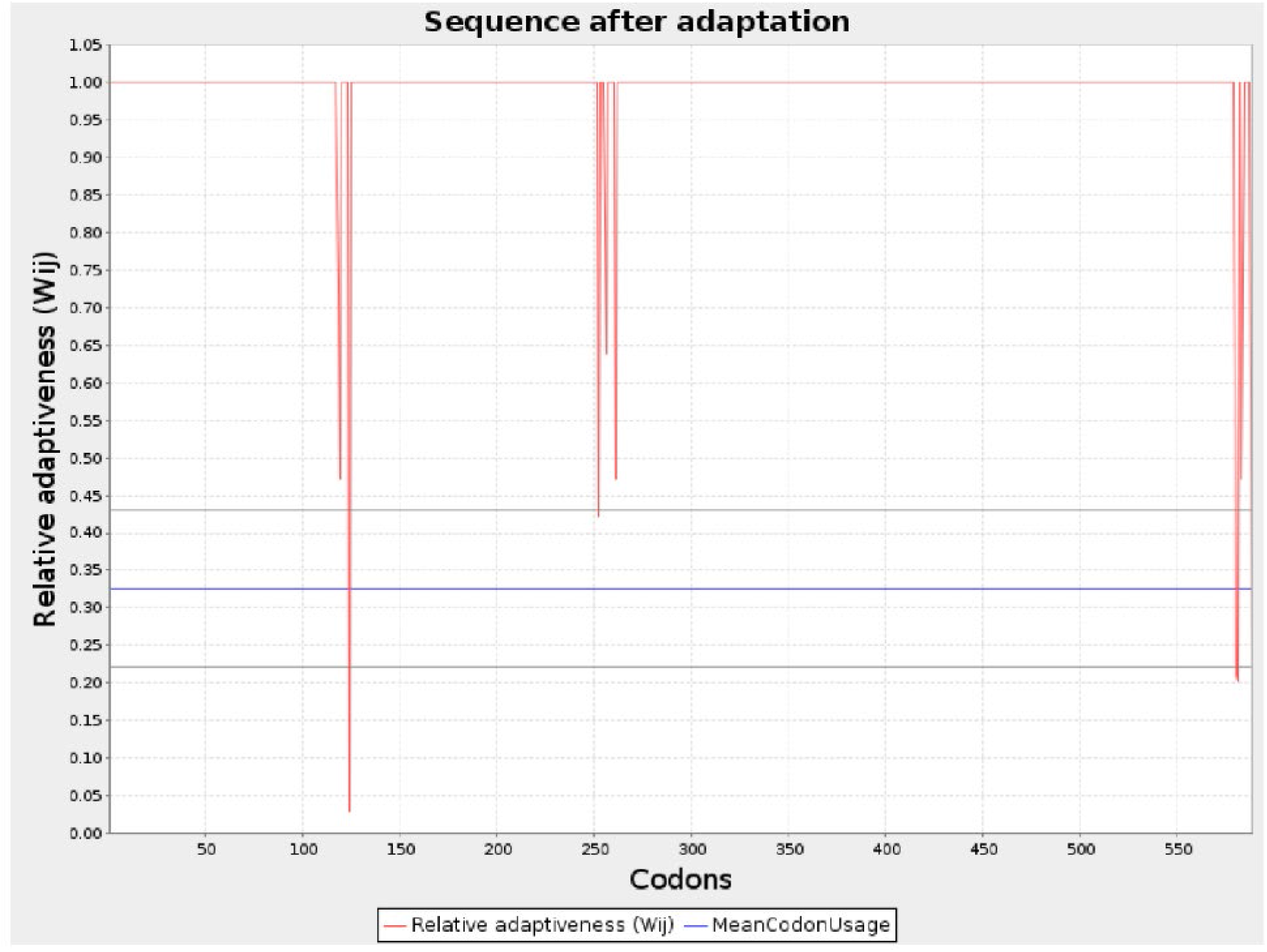
*In silico* codon optimization profile of the chimeric vaccine construct (MPVX-3) for heterologous expression in *Escherichia coli* (strain K12). Relative adaptiveness (*w*ᵢⱼ) along the 580-codon sequence following adaptation using the JCat (Java Codon Adaptation Tool) server (red line), benchmarked against the mean codon usage threshold (blue line). Codons scoring *wᵢⱼ* = 1.0 correspond to the most frequently used synonymous codons in the host expression system. Localized dips indicate regions constrained to eliminate unwanted restriction sites, GC extreme regions, or destabilizing mRNA secondary elements.

### Recurrent neural network-guided optimization of mRNA secondary structure and translation initiation kinetics

To ensure that high codon adaptation translated into unimpeded protein expression in host translation systems, the synthetic transcript encoding MPXV-3 was modeled through a recurrent neural network (RNN)-based generative optimization framework, RiboDecode, balancing the translation initiation rate against global conformational stability (Figure 8C, D). Analysis of the 5’ untranslated and leader initiation window (nucleotides 1–60) revealed a discrete, low-energy secondary folding threshold (ΔG = −11.6 kcal mol^-1^) characterized by an unconstrained single-stranded architecture across the first 18 nucleotides (Figure 8C). This single-stranded 5’ architecture preserves steric accessibility for the pre-initiation ribosomal scanning machinery, avoiding premature scanning arrest. Across the full-length 1,780 nt open reading frame, the transcript adopted a dense, highly stable global thermodynamic architecture with a minimum free energy (MFE) of ΔG = − 678.1 kcal mol^-1^ (Figure 8D). Sliding-window base-pair density profiling demonstrated a homeostatic distribution fluctuating between 0.35 and 0.82 across the coding core, devoid of persistent hyper-dense secondary hairpins (>0.95 density) that induce ribosomal pausing [104]. Together, the juxtaposition of an open, low-affinity initiation terminal with a thermodynamically stabilized coding body demonstrates that deep learning-guided sequence optimization preserves translation initiation potential while maximizing cellular transcript half-life and preventing steric ribosomal collisions [12], [105].

**Figure 8.**
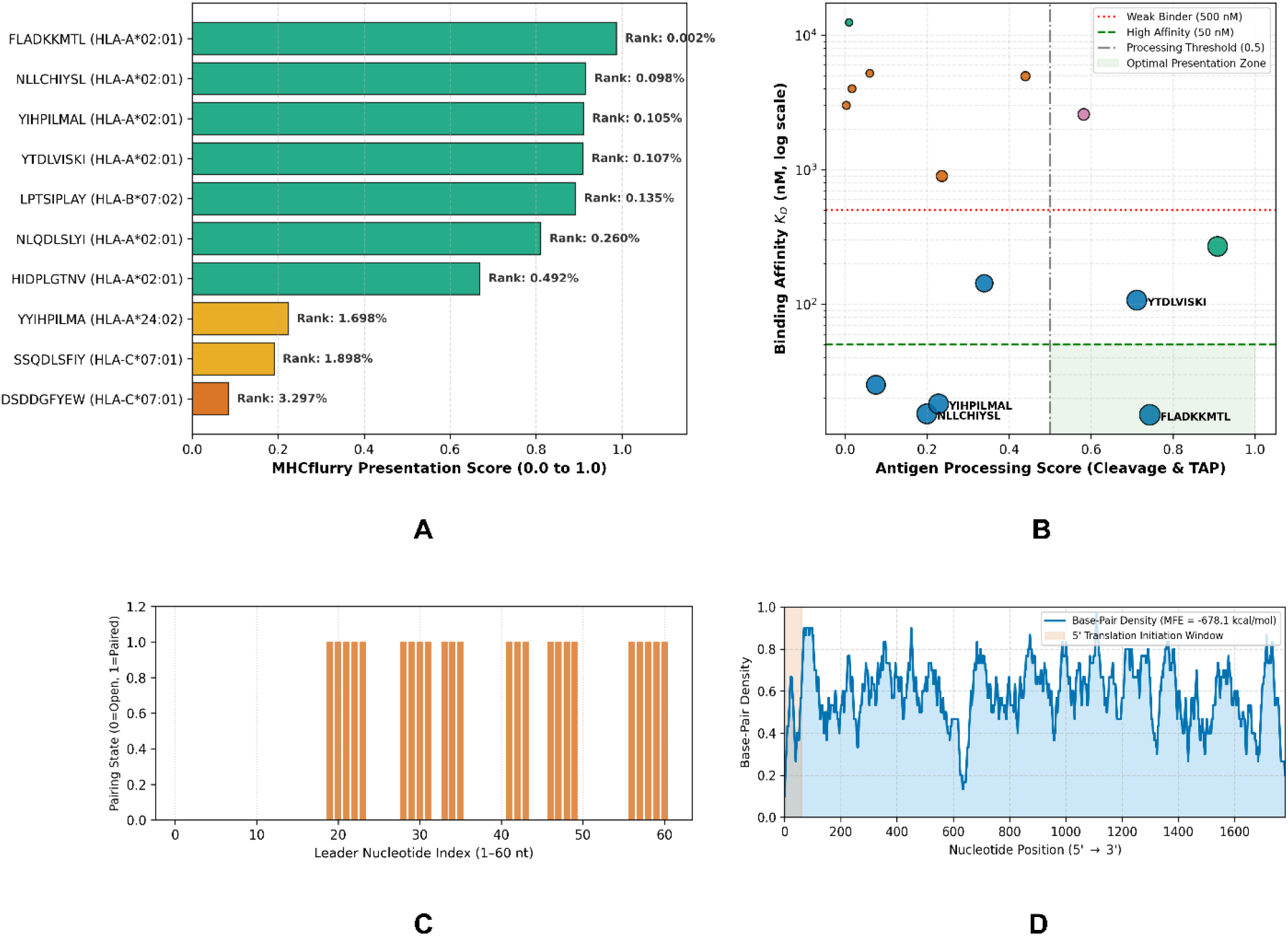
Multidimensional deep learning profiling of antigen processing, human leukocyte antigen (HLA) presentation, and mRNA thermodynamic stability for the lead Mpox vaccine construct. (A) MHCflurry 2.0 presentation ranking across representative human HLA Class I alleles. Bars indicate composite presentation scores (0.0–1.0), color-coded by presentation percentile rank: green (≤0.5%, best), yellow (0.5%–2.0%, moderate), and orange (>2.0%, low). Annotated values highlight top HLA-A*02:01-restricted candidates FLADKKMTL(0.002%) and NLLCHIYSL (0.098%). (B) Dual-parameter profiling of intracellular processing against HLA groove binding affinity (*K_D_*, nM; log scale). Reference lines denote weak binding (500nM, red dotted), high affinity (50nM, green dashed), and efficient processing (≥0.50, grey dash-dot). The green quadrant marks the optimal high-affinity processing space; marker size scales with overall presentation score. (C) Base-pairing profile across the 5’ mRNA leader sequence (nt 1–60). Binary pairing states (0 = open, 1 = paired) show an unconstrained single-stranded initiation window (ΔG = –11.6 kcal mol^-1^) through the first 18 nucleotides, supporting ribosomal assembly. (D) Secondary structure profile of the full 1,780 nt mRNA transcript modeled via ViennaRNA. The base-pairing density curve reflects a thermodynamically favorable architecture (ΔG = –678.1 kcal mol^-1^) without hyper-stable local hairpins (>0.95 density), supporting steady elongation and transcript stability. The shaded box marks the 5’ leader window shown in Panel C.

**Figure 9.**
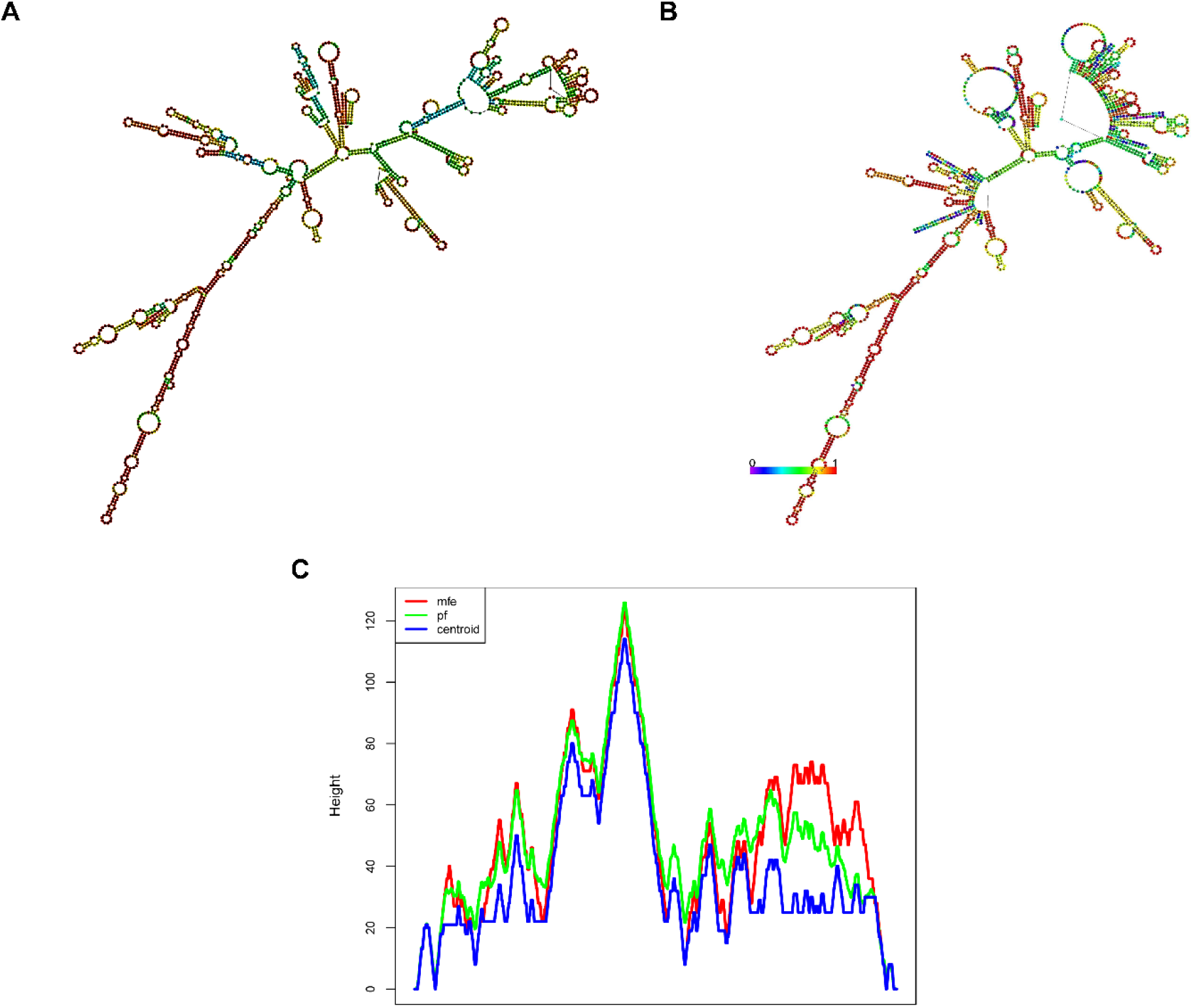
Secondary structure modeling and thermodynamic stability profiling of the codon-optimized MVX-3 mRNA construct. (a) Minimum free energy (MFE) secondary structure prediction of the 1,777-nt transcript calculated via RNAfold (ΔG = –676.10 kcal mol^-1^, demonstrating dense base-pairing across structured hairpin stems. (b) Thermodynamic ensemble centroid secondary structure (ΔG = –543.62 kcal mol^-1^, ensemble free energy ΔG = –700.30 kcal mol^-1^, color-annotated by base-pairing probability spectrum (0 to 1, violet to red). (c) Mountain plot representation illustrating the height-based structural landscape along the sequence coordinates for the MFE structure (red), thermodynamic partition function (pf, green), and centroid conformation (blue), reflecting conformational congruency across central multi-branch loop domains.

A central failure mode in the translational deployment of synthetic mRNA chimeras is the kinetic antagonism between *in vivo* transcript stability and ribosomal processivity: maximizing thermodynamic base-pairing often stabilizes the molecule against intracellular ribonucleases at the expense of creating secondary roadblocks that prompt ribosome stalling and translation abortion [105], [106]. Computational algorithms addressing this trade-off, such as LinearDesign, have demonstrated that co-optimizing minimum free energy alongside codon adaptation significantly amplifies protein expression and half-life by suppressing spontaneous RNA hydrolysis [12]. Our RNN-guided transcript profile for MPXV-3 confirms this cooperative dynamic, achieving a negative free energy threshold (ΔG = − 678.1 kcal mol^-1^) without forming the severe local folding traps that trigger translational traffic jams [104]. Crucially, the functional viability of this architecture depends on the structural polarization resolved in Figure 8C, D: while the elongation body remains tightly base-paired, the initiation portal is deliberately de-structured (ΔG = −11.6 kcal mol^-1^), leaving nucleotides 1–18 in an open conformation. This spatial configuration aligns with experimental evidence that weakly folded 5’ leaders are essential for rapid 40S subunit recruitment and unhindered scanning of the initiation complex [107]. However, translating these *in silico* predictions to wet-lab systems involves biological variables; while our moving-average density profiles show no hyper-stable hairpins, high-density bacterial expression (e.g., under a T7 promoter in pET-28a(+)) can slow local elongation if transient secondary-folding knots interact synergistically with minor tRNA pool fluctuations [105], [106]. Consequently, cell-free translation assays (e.g., rabbit reticulocyte lysate or HeLa ribosome profiling) remain the necessary empirical bridge to ensure that the predicted kinetic accessibility translates into high-titer production of the untruncated 588-AA polypeptide.

Secondary structure profiling of the codon-optimized MVX-3 transcript (1,777 nt) via ViennaRNA RNAfold revealed favorable thermodynamic stability, yielding a minimum free energy (MFE) of –676.10 kcal mol^-1^ and a thermodynamic ensemble free energy of –700.30 kcal mol^-1^ with a centroid distance of –543.62 kcal mol^-1^ (Figure 5A and 5B). Mountain plot representations demonstrated close alignment between the MFE trajectory, the partition function, and the centroid conformation across the sequence coordinates, confirming conserved secondary structure architectures across the ensemble (Figure 5C).

The substantial negative folding free energy (ΔG = –676.10 kcal mol^-1^, –0.38 kcal mol^-1^) indicates that the engineered MVX-3 mRNA forms a stable secondary fold resistant to spontaneous ribonuclease-mediated degradation. Furthermore, the decentralized multi-branch loop topology observed across the thermodynamic ensemble facilitates localized helicase unwinding during ribosomal elongation, supporting both transcript longevity and translational competence.

## Conclusion

This study demonstrates the transition from traditional, sequence-reliant reverse vaccinology to a deep learning-augmented modern framework for vaccine structural design. By targeting the most virulent components of the Mpox virus proteome, we engineered three next-generation multi-epitope subunit vaccines designed to induce robust, multi-antigenic immunity and minimize the risk of mutational viral escape. By integrating a high-fidelity artificial intelligence validation layer for our final MVX-3 candidate, we addressed the classic false-positive bottlenecks that have historically limited *in silico* workflows. Applying ESM-2 protein language models confirmed that our chimeric constructs maintain native-like structural stability across synthetic linker junctions, significantly reducing the likelihood of proteolysis. Furthermore, utilizing structure-informed graph neural networks (GNNs) allowed us to model the dynamic, non-linear physical interactions of the immune synapse, ensuring high-confidence binding and presentation across diverse human HLA alleles. Finally, RNN-driven mRNA kinetic optimization validated that these constructs are optimized for high-yield expression, directly bridging the gap between computational discovery and biopharmaceutical manufacturing. Ultimately, this multi-objective computational pipeline provides a highly rigorous, scalable blueprint for vaccine design. By validating structural naturalness, immunogenicity, and translational kinetics prior to *in vitro* synthesis, this framework sets a new standard for proactive, rapid-response countermeasure development against emerging orthopoxvirus threats.

## Supporting information

https://doi.org/10.5281/zenodo.22865532

## Acknowledgement

The authors acknowledge the Bioinformatics Laboratory (Biolab), Bangladesh, for their molecular dynamics (MD) simulation support.

## Supporting information

All supporting information will be available in the supplementary files section.

## Code and data availability

All source code is available on GitHub (https://github.com/aakilkhanemon/AI-assisted-vaccine-design). All associated data, including deep learning immunopeptidomics screening, and RNN-driven mRNA kinetic optimization, are available at Zenodo (https://doi.org/10.5281/zenodo.22865532).

