## Supplementary material for "AI-Driven Design of Next-Generation Immunoinformatics Multi-Epitope Subunit Vaccine Targeting the Most Virulent Mpox Proteome: A Structural and Kinetic Validation Framework": https://doi.org/10.5281/zenodo.22865532

**Contents**

Supplementary Methods……………………………………………………………………………….. 1-2
Supplementary Tables……………………………………………………………...………….......... 2-9
Supplementary Figures……………………………………………………………………………….. 9-11

### Supplementary Computational Methods

#### Text S1. GNN-based modeling of the immune synapse

To quantitatively evaluate binding engagement between candidate peptide epitopes and the human T-cell receptor (TCR)–major histocompatibility complex (MHC) interface, structural models of the ternary complex (peptide–HLA-A*11:01–TCR) were converted into molecular graph representations. Each amino acid was represented as an individual node centered at its *Cα* spatial coordinate. Spatial adjacency edges were established between pairs of residues within a 10.0 Å Euclidean distance cutoff. Inter-chain contact edges were weighted inversely using AlphaFold Predicted Aligned Error (PAE) matrices to down-weight uncertain interface conformations. Node feature vectors encoded amino acid physicochemical properties, solvent accessibility, and sequence-derived transformer embeddings. Message passing across the graph used a structure-informed graph neural network (GNN) architecture with multi-head edge-attention mechanisms. Normalized attention weights ($\alpha_{ij}$) between the 9-mer epitope residues and the TCR CDR3 loops (α and β) were extracted to map functional contact drivers at residue-level resolution. The final binding probability ($p_{\text{binding}}$) was computed via global graph pooling followed by a multilayer perceptron (MLP) with a terminal sigmoid activation function, where $p \geq 0.50$ served as the canonical classification threshold for cognate TCR recognition.

#### Text S2. Latent space validation

To verify that artificial multi-epitope junctions (incorporating EAAAK, AAY, and GPGPG linkers) did not introduce structural instability or unphysical sequences, chimeras were evaluated using the evolutionary-scale protein language model ESM-2 (esm2_t33_650M_UR50D, 650M parameters). We quantified sequence naturalness and structural harmony using a pseudo-masked log-likelihood estimation protocol. For an engineered construct of length N, each residue position *xi* was iteratively replaced with the designated mask token, while leaving flanking contextual residues unperturbed. We passed the masked representation through ESM-2 and recorded the conditional Softmax probability $P(x_i|x_{-i})$ corresponding to the native amino acid. Sequence-wide pseudo-perplexity (PPL) was calculated as:

$$\mathrm{PPL} = \exp\left(-\frac{1}{N}\sum_{i=1}^{N}\ln P\left(x_i \mid x_{\setminus i}\right)\right)$$

Constructs maintaining pseudo-perplexity scores within the baseline regime of natural globular proteins PPL ≤ 12.0 were deemed structurally viable, ensuring low conformational strain as well as minimal risk of aggregation or proteolysis.

**Text S3. Deep learning-augmented immunopeptidomics and antigen processing profiling**

Physiological antigen presentation was modeled using MHCflurry v2.0 (models_class1_presentation), combining neural network predictors for peptide MHC binding affinity with deep models of antigen processing. Candidate epitope sequences were evaluated across high-frequency human HLA Class I reference alleles (eg, HLA-A*02:01, HLA-A*24:02, HLA-B*07:02, HLA-B*08:01, HLA-C*07:01, and HLA-A*11:01). For each peptide–allele pair, the pipeline computed:

- An affinity prediction quantifying the equilibrium dissociation constant ($K_D$ in nM).
- An antigen processing score derived from proteasomal cleavage and TAP transport heuristics.
- An integrated presentation score with its corresponding presentation percentile rank calibrated against reference human proteome decoys. Epitopes achieving a presentation percentile rank <2.0% were classified as presentation-competent targets.

**Text S4. Recurrent neural network-guided optimization of mRNA secondary structure and translation initiation kinetics**

To maximize heterologous expression and prevent premature ribosomal termination during therapeutic mRNA delivery, coding sequences were optimized via a recurrent neural network (RNN) driven framework paired with thermodynamic folding models. The wild-type amino acid sequence was reverse-translated while simultaneously optimizing the codon adaptation index (CAI) for human host expression and avoiding unfavorable GC-content extremes (targeted between 45% and 65%). Codon selection was evaluated using bidirectional long short-term memory (BiLSTM) networks trained on human translation initiation and elongation kinetics. Downstream mRNA folding landscapes were modelled using LinearFold and ViennaRNA packages to compute global minimum free energy (ΔG_MFE, in kcal $mol^{-1}$) and structural accessibility around the 5' untranslated region (UTR) translation initiation site (Kozak consensus). Candidate transcripts were filtered to ensure low propensity for stable inhibitory hairpins near the initiation complex while maintaining sufficient secondary structure within the open reading frame (ORF) to optimize transcript half-life and cellular persistence.

### Supplementary Tables

**Table S1.** Selected Mpox virus (MPXV) target proteins and corresponding NCBI GenBank accession identifiers utilized for epitope prioritization and construct assembly. Functional annotation and reference sequence accession codes for structural and non-structural viral proteins analyzed in this study.

| Serial number | Protein name | Accession number |
|---|---|---|
| **1** | Intracellular viral protein | NP_536598.1 |
| **2** | Viral membrane formation protein | NP_536549.1 |
| **3** | Virion protein D2 | NP_536526.1 |
| **4** | Virion core protein D3 | WQK80685.1 |
| **5** | Membrane protein CL5 | NP_536511.1 |
| **6** | Assembly protein G7 | NP_536504.1 |
| **7** | Virion core protein P4b | NP_536541.1 |
| **8** | Viral core cysteine proteinase | UVT84927.1 |

**Table S2.** *In silico* immunogenic profiling, allergenicity classification, and multi-parametric safety screening of candidate Mpox virus (MPXV) proteins. Quantitative antigenicity scores determined via VaxiJen (threshold

≥0.45), allergenicity categorization via AllerTOP/AllergenFP, and toxicity predictions via ToxinPred across the eight initial viral targets. Biophysical/physicochemical benchmarking evaluated thermodynamic stability via Expasy ProtParam; proteins with an instability index >40.0 were classified as structurally unstable (Fail). Only virion core protein P4b and viral core cysteine proteinase satisfied all cumulative criteria (antigenicity ≥0.45, non-allergen, non-toxin, and instability index <40.0) and were prioritized for downstream epitope mapping.

| Protein name | Antigenicity | Allergenicity | Toxicity | Biophysical/ physicochemical benchmark |
|---|---|---|---|---|
| Intracellular viral protein | 0.36 | Allergen | Non-toxin | Fail |
| Viral membrane formation protein | 0.46 | Non-allergen | Non-toxin | Fail |
| Virion protein D2 | 0.39 | Non-allergen | Non-toxin | Fail |
| Virion core protein D3 | 0.39 | Non-allergen | Non-toxin | Fail |
| Membrane protein CL5 | 0.45 | Non-allergen | Non-toxin | Fail |
| Assembly protein G7 | 0.35 | Non-allergen | Non-toxin | Fail |
| **Virion core protein P4b** | 0.45 | Non-allergen | Non-toxin | Pass |
| **Viral core cysteine proteinase** | 0.52 | Non-allergen | Non-toxin | Pass |

**Table S3.** Biophysical and physicochemical characterization of the prioritized MPXV target proteins. Primary sequence parameters calculated via the ExPASy ProtParam server, including molecular weight (MW in Da), theoretical isoelectric point (pI), extinction coefficient ($M^{-1}$ $cm^{-1}$ at 280 nm), predicted mammalian *in vitro* half-life, aliphatic index, instability index, and grand average of hydropathicity (GRAVY). Both vaccine candidates demonstrate thermal stability (aliphatic index >80), classified structural stability in solution (instability index <40.0), and hydrophilic character (GRAVY <0).

| Name of protein sequence | Total amino acid | MW | Theoretical pI | Ext. coefficient (in $M^{-1}$ $cm^{-1}$) | Est. half-life (in mammalian cell) hours | Aliphatic index | Instability index | GRAVY |
|---|---|---|---|---|---|---|---|---|
| **Virion core protein P4b** | 644 | 68994.7 | 6.22 | 58970 | 30 | 85.2 | 39.0 | −0.23 |
| **Viral core cysteine proteinase** | 423 | 49023.57 | 7.85 | 45560 | 30 | 83.14 | 33.8 | −0.22 |

**Table S4.** *In silico* screening, binding affinity prioritization, and immunogenic profiling of MHC Class I epitopes derived from Mpox virus (MPXV) virion core protein P4b. Epitope prediction and percentile score ranking were performed via the IEDB analysis resource (NetMHCpan/consensus algorithm), followed by binary screening for antigenicity (VaxiJen), allergenicity (AllerTOP), toxicity (ToxinPred), and cross-reactivity with the human proteome (NCBI BLASTp). Bolded sequences denote candidates prioritized based on top percentile affinity and designated criteria.

| Epitope | Start | End | Percentile score | Antigenicity | Allergenicity | Toxicity | Human homology |
|---|---|---|---|---|---|---|---|
| **FTENGPWMY** | 536 | 544 | 0.995 | Antigenic | Non-allergen | Non-toxin | Non-homologous |
| **SSQDLSFIY** | 215 | 223 | 0.889 | Antigenic | Non-allergen | Non-toxin | Non-homologous |
| SMDSDDGFY | 571 | 579 | 0.862 | Non-antigenic | Non-allergen | Non-toxin | Non-homologous |
| **YIHPILMAL** | 231 | 239 | 0.843 | Antigenic | Non-allergen | Non-toxin | Non-homologous |
| AVDPHLMKI | 328 | 336 | 0.823 | Non-antigenic | Non-allergen | Non-toxin | Non-homologous |
| MLNGIMYRL | 498 | 506 | 0.808 | Non-antigenic | Non-allergen | Non-toxin | Non-homologous |
| **TLNNASVSV** | 155 | 163 | 0.761 | Antigenic | Non-allergen | Non-toxin | Non-homologous |
| ILASILSIV | 351 | 359 | 0.738 | Non-antigenic | Non-allergen | Non-toxin | Non-homologous |
| RLTEDNPIV | 280 | 288 | 0.726 | Non-antigenic | Allergen | Non-toxin | Non-homologous |
| YVHGDTYSL | 251 | 259 | 0.635 | Non-antigenic | Allergen | Non-toxin | Non-homologous |
| **KLQAAPYIV** | 380 | 388 | 0.634 | Antigenic | Non-allergen | Non-toxin | Non-homologous |
| MTDEEQILA | 345 | 353 | 0.547 | Non-antigenic | Allergen | Non-toxin | Non-homologous |

| | | | | | | | |
|---|---|---|---|---|---|---|---|
| IVNGAMMFY | 456 | 464 | 0.512 | Non-antigenic | Allergen | Non-toxin | Non-homologous |
| ALFGIKLPA | 238 | 246 | 0.501 | Non-antigenic | Non-allergen | Non-toxin | Non-homologous |
| **HIDPLGTNV** | 438 | 446 | 0.496 | Antigenic | Non-allergen | Non-toxin | Non-homologous |
| ELFQLLSHV | 144 | 152 | 0.484 | Non-antigenic | Allergen | Non-toxin | Non-homologous |
| **LIDDTPIDV** | 485 | 493 | 0.483 | Antigenic | Non-allergen | Non-toxin | Non-homologous |
| FLKGHYTIL | 527 | 535 | 0.381 | Non-antigenic | Allergen | Non-toxin | Non-homologous |
| **DSDDGFYEW** | 573 | 581 | 0.331 | Antigenic | Non-allergen | Non-toxin | Non-homologous |
| **LLSHVKSTL** | 148 | 156 | 0.327 | Antigenic | Non-allergen | Non-toxin | Non-homologous |

**Table S5.** *In silico* screening, binding affinity prioritization, and immunogenic profiling of MHC Class II epitopes derived from Mpox virus (MPXV) virion core protein P4b. Epitope prediction and percentile score ranking were performed via the IEDB analysis resource (NetMHCpan/consensus algorithm), followed by binary screening for antigenicity (VaxiJen), allergenicity (AllerTOP), toxicity (ToxinPred), and cross-reactivity with the human proteome (NCBI BLASTp). Bolded sequences denote candidates prioritized based on top percentile affinity and designated criteria.

| Epitope | Start | End | Percentile score | Antigenicity | Allergenicity | Toxicity | Human homology |
|---|---|---|---|---|---|---|---|
| **PILMALFGIKLPALE** | 234 | 248 | 1.10 | Antigenic | Non-allergen | Non-toxin | Non-homologous |
| **HPILMALFGIKLPAL** | 233 | 247 | 1.30 | Antigenic | Non-allergen | Non-toxin | Non-homologous |
| NELFQLLSHVKSTLN | 143 | 157 | 1.60 | Non-antigenic | Allergen | Non-toxin | Non-homologous |
| RNELFQLLSHVKSTL | 142 | 156 | 1.60 | Non-antigenic | Allergen | Non-toxin | Non-homologous |
| **YYIHPILMALFGIKL** | 230 | 244 | 1.60 | Antigenic | Non-allergen | Non-toxin | Non-homologous |
| ELFQLLSHVKSTLNN | 144 | 158 | 1.80 | Non-antigenic | Allergen | Non-toxin | Non-homologous |
| **YIHPILMALFGIKLP** | 231 | 245 | 1.80 | Antigenic | Non-allergen | Non-toxin | Non-homologous |
| **IHPILMALFGIKLPA** | 232 | 246 | 2.00 | Antigenic | Non-allergen | Non-toxin | Non-homologous |
| **NRNELFQLLSHVKST** | 141 | 155 | 2.00 | Antigenic | Non-allergen | Non-toxin | Non-homologous |
| KSYNYMLLVNRLTED | 270 | 284 | 2.10 | Non-antigenic | Non-allergen | Non-toxin | Non-homologous |
| **VKSYNYMLLVNRLTE** | 269 | 283 | 2.10 | Antigenic | Non-allergen | Non-toxin | Non-homologous |
| SYNYMLLVNRLTEDN | 271 | 285 | 2.40 | Non-antigenic | Non-allergen | Non-toxin | Non-homologous |
| **RVKSYNYMLLVNRLT** | 268 | 282 | 2.80 | Antigenic | Non-allergen | Non-toxin | Non-homologous |
| **ILSIVGFRPTLVSVA** | 355 | 369 | 2.90 | Antigenic | Non-allergen | Non-toxin | Non-homologous |
| **ILMALFGIKLPALEN** | 235 | 249 | 3.10 | Antigenic | Non-allergen | Non-toxin | Non-homologous |
| **LMALFGIKLPALENA** | 236 | 250 | 3.10 | Antigenic | Non-allergen | Non-toxin | Non-homologous |
| **LSIVGFRPTLVSVAR** | 356 | 370 | 3.10 | Antigenic | Non-allergen | Non-toxin | Non-homologous |
| YNYMLLVNRLTEDNP | 272 | 286 | 4.30 | Non-antigenic | Allergen | Non-toxin | Non-homologous |
| TNRNELFQLLSHVKS | 140 | 154 | 4.40 | Non-antigenic | Allergen | Non-toxin | Non-homologous |
| **THPLVLIHSRASPRI** | 166 | 180 | 4.90 | Antigenic | Non-allergen | Non-toxin | Non-homologous |

**Table S6.** *In silico* mapping, safety profiling, and prioritization of linear B-cell epitopes from Mpox virus (MPXV) virion core protein P4b. Linear B-cell epitopes identified via the IEDB server, characterized for antigenicity (VaxiJen), allergenicity (AllerTOP), toxicity (ToxinPred), and cross-reactivity with the human proteome (NCBI BLASTp). Bolded sequences denote candidates satisfying all benchmark criteria (antigenic, non-allergen, non-toxin, and non-homologous to human proteins) prioritized for vaccine construct integration.

| Epitope | Start | End | Antigenicity | Allergenicity | Toxicity | Human homology |
|---|---|---|---|---|---|---|
| **AGLKSSYTNQTLSLVDEDHIHTSDKS** | 15 | 40 | Antigenic | Non-allergen | Non-toxin | Non-homologous |
| **GARNQRTKPKRTGNDQSQQPIKKDCMVSIDEVASTHDWSTR** | 61 | 101 | Antigenic | Non-allergen | Non-toxin | Non-homologous |
| KYDTSNF | 117 | 123 | Non-antigenic | Allergen | Non-toxin | Non-homologous |
| NIVRTNRNEL | 136 | 145 | Non-antigenic | Non-allergen | Non-toxin | Non-homologous |
| KELDKIYSPSNH | 185 | 196 | Non-antigenic | Non-allergen | Non-toxin | Non-homologous |
| **FTDMSSSQDLSFIYRKPETN** | 210 | 229 | Antigenic | Non-allergen | Non-toxin | Non-homologous |
| YEFRRVKS | 264 | 271 | Non-antigenic | Non-allergen | Non-toxin | Non-homologous |
| CSQVMTDE | 341 | 348 | Non-antigenic | Allergen | Non-toxin | Non-homologous |
| PMNGISYD | 371 | 378 | Non-antigenic | Non-allergen | Non-toxin | Non-homologous |
| KMITTSDSP | 393 | 401 | Non-antigenic | Allergen | Non-toxin | Non-homologous |
| GYGRCSGVTHIDPLGTNVMGS | 429 | 449 | Non-antigenic | Allergen | Non-toxin | Non-homologous |
| **QNKNMFGG** | 469 | 476 | Antigenic | Non-allergen | Non-toxin | Non-homologous |
| YTGFRSLIDDTPIDV | 479 | 493 | Non-antigenic | Non-allergen | Non-toxin | Non-homologous |
| GDQFFDCGS | 515 | 523 | Non-antigen | Non-allergen | Non-toxin | Non-homologous |
| LSVFNPGARNA | 547 | 557 | Non-antigenic | Non-allergen | Non-toxin | Non-homologous |
| **YKKLSMDSDDG** | 567 | 577 | Antigenic | Non-allergen | Non-toxin | Non-homologous |
| YEWLNGDGSVFAASKQQMLMNH | 579 | 600 | Non-antigenic | Allergen | Non-toxin | Non-homologous |
| ANFDDDLLTM | 602 | 611 | Non-antigenic | Allergen | Non-toxin | Non-homologous |
| **DQYISARHI** | 632 | 640 | Antigenic | Non-allergen | Non-toxin | Non-homologous |

**Table S7.** *In silico* identification, binding affinity ranking, and multi-parametric safety profiling of MHC Class I epitopes derived from Mpox virus (MPXV) viral core cysteine proteinase. Epitope mapping and percentile score ranking were conducted via the IEDB analysis resource, followed by evaluation for antigenicity (VaxiJen), allergenicity (AllerTOP), toxicity (ToxinPred), and cross-reactivity with the human proteome (NCBI BLASTp). Bolded sequences denote candidates that met all benchmark criteria (antigenic, non-allergen, non-toxin, and non-homologous to human self-antigens) prioritized for downstream construct integration.

| Epitope | Start | End | Percentile score | Antigenicity | Allergenicity | Toxicity | Human homology |
|---|---|---|---|---|---|---|---|
| LTDELTTRY | 92 | 100 | 0.998 | Non-antigenic | Non-allergen | Non-toxin | Non-homologous |
| **FLADKKMTL** | 359 | 367 | 0.982 | Antigenic | Non-allergen | Non-toxin | Non-homologous |
| LVISKIPEL | 7 | 15 | 0.846 | Non-antigenic | Non-allergen | Non-toxin | Non-homologous |
| FMNKHILSY | 210 | 218 | 0.792 | Non-antigenic | Allergen | Non-toxin | Non-homologous |
| **NLQDLSLYI** | 374 | 382 | 0.719 | Antigenic | Non-allergen | Non-toxin | Non-homologous |
| **NLLCHIYSL** | 19 | 27 | 0.674 | Antigenic | Non-allergen | Non-toxin | Non-homologous |
| FTNLLCHIY | 17 | 25 | 0.669 | Non-antigenic | Allergen | Non-toxin | Non-homologous |
| **LTTRYHSIY** | 96 | 104 | 0.456 | Antigenic | Non-allergen | Non-toxin | Non-homologous |
| FSDGFNTNH | 278 | 286 | 0.451 | Non-antigenic | Allergen | Non-toxin | Non-homologous |
| **SQFAYGSEV** | 179 | 187 | 0.436 | Antigenic | Non-allergen | Non-toxin | Non-homologous |
| **LADKKMTLF** | 360 | 368 | 0.402 | Antigenic | Non-allergen | Non-toxin | Non-homologous |
| **YTDLVISKI** | 4 | 14 | 0.372 | Antigenic | Non-allergen | Non-toxin | Non-homologous |
| **FAYGSEVDY** | 181 | 189 | 0.275 | Antigenic | Non-allergen | Non-toxin | Non-homologous |
| NIPTEFHHY | 262 | 270 | 0.269 | Non-antigenic | Allergen | Non-toxin | Non-homologous |
| **SSDELDQKK** | 82 | 90 | 0.246 | Antigenic | Non-allergen | Non-toxin | Non-homologous |
| **LPTSIPLAY** | 109 | 117 | 0.229 | Antigenic | Non-allergen | Non-toxin | Non-homologous |
| KLTTKLNRI | 410 | 418 | 0.224 | Non-antigenic | Non-allergen | Non-toxin | Non-homologous |
| SLAGLCSNI | 26 | 34 | 0.216 | Non-antigenic | Allergen | Non-toxin | Non-homologous |
| **VIYDKKQCL** | 245 | 253 | 0.203 | Antigenic | Non-allergen | Non-toxin | Non-homologous |
| FLTNCNGYV | 39 | 47 | 0.189 | Non-antigenic | Allergen | Non-toxin | Non-homologous |

**Table S8.** *In silico* identification, binding affinity ranking, and multi-parametric safety profiling of MHC Class II epitopes derived from Mpox virus (MPXV) viral core cysteine proteinase. Epitope mapping and percentile score ranking were conducted via the IEDB analysis resource, followed by evaluation for antigenicity (VaxiJen), allergenicity (AllerTOP), toxicity (ToxinPred), and cross-reactivity with the human proteome (NCBI BLASTp). Bolded sequences denote candidates that met all benchmark criteria (antigenic, non-allergen, non-toxin, and non-homologous to human self-antigens) prioritized for downstream construct integration.

| Epitope | Start | End | Percentile score | Antigenicity | Allergenicity | Toxicity | Human homology |
|---|---|---|---|---|---|---|---|
| **FDMRFLNSLAIHEKF** | 193 | 207 | 3.30 | Antigenic | Non-allergen | Non-toxin | Non-homologous |

| | | | | | | | |
|---|---|---|---|---|---|---|---|
| **QFDMRFLNSLAIHEK** | 192 | 206 | 3.30 | Antigenic | Non-allergen | Non-toxin | Non-homologous |
| KSFKSLKKVYTFFKF | 345 | 359 | 3.70 | Non-antigenic | Non-allergen | Non-toxin | Non-homologous |
| NRSIKNLVSQFAYGS | 171 | 185 | 3.70 | Non-antigen | Non-allergen | Non-toxin | Non-homologous |
| PKSFKSLKKVYTFFK | 344 | 358 | 3.70 | Non-antigenic | Non-allergen | Non-toxin | Non-homologous |
| RSIKNLVSQFAYGSE | 172 | 186 | 3.70 | Non-antigenic | Allergen | Non-toxin | Non-homologous |
| PPKSFKSLKKVYTFF | 343 | 357 | 3.90 | Non-antigenic | Non-allergen | Non-toxin | Non-homologous |
| **GQFDMRFLNSLAIHE** | 191 | 205 | 4.10 | Antigenic | Non-allergen | Non-toxin | Non-homologous |
| TPPKSFKSLKKVYTF | 342 | 356 | 4.80 | Non-antigenic | Non-allergen | Non-toxin | Non-homologous |
| YTFFKFLADKKMTLF | 354 | 368 | 6.30 | Non-antigenic | Non-allergen | Non-toxin | Non-homologous |
| SIKNLVSQFAYGSEV | 173 | 187 | 6.50 | Non-antigenic | Allergen | Non-toxin | Non-homologous |
| TFFKFLADKKMTLFK | 355 | 369 | 6.50 | Non-antigenic | Non-allergen | Non-toxin | Non-homologous |
| VYTFFKFLADKKMTL | 353 | 367 | 6.50 | Non-antigenic | Non-allergen | Non-toxin | Non-homologous |
| SNRSIKNLVSQFAYG | 170 | 184 | 6.70 | Non-antigenic | Non-allergen | Non-toxin | Non-homologous |
| MSNRSIKNLVSQFAY | 169 | 183 | 6.80 | Non-antigenic | Non-allergen | Non-toxin | Non-homologous |
| IKNLVSQFAYGSEVD | 174 | 188 | 7.30 | Non-antigenic | Allergen | Non-toxin | Non-homologous |
| **IGQFDMRFLNSLAIH** | 190 | 204 | 7.70 | Antigenic | Non-allergen | Non-toxin | Non-homologous |
| KVYTFFKFLADKKMT | 352 | 366 | 8.70 | Non-antigenic | Allergen | Non-toxin | Non-homologous |
| LLCHIYSLAGLCSNI | 20 | 34 | 8.90 | Non-antigenic | Non-allergen | Non-toxin | Non-homologous |
| **LCHIYSLAGLCSNID** | 21 | 35 | 9.00 | Antigenic | Non-allergen | Non-toxin | Non-homologous |

**Table S9.** *In silico* mapping, safety evaluation, and prioritization of linear B-cell epitopes from Mpox virus (MPXV) viral core cysteine proteinase. Linear B-cell determinants identified via IEDB and evaluated for antigenicity (VaxiJen), allergenicity (AllerTOP), toxicity (ToxinPred), and human proteome cross-reactivity (NCBI BLASTp). Bolded sequences indicate prioritized candidates evaluated against downstream construct inclusion criteria.

| Epitope | Start | End | Antigenicity | Allergenicity | Toxicity | Human homology |
|---|---|---|---|---|---|---|
| SKFLTNCN | 37 | 44 | Non-antigenic | Non-allergen | Non-toxin | Non-homologous |
| VEKYDKSTTA | 48 | 57 | Non-antigenic | Allergen | Non-toxin | Non-homologous |
| GHLSRPNSSDELDQKKEL | 75 | 92 | Non-antigenic | Allergen | Non-toxin | Non-homologous |
| TTRYHSIYDVFELP | 97 | 110 | Non-antigenic | Allergen | Non-toxin | Non-homologous |
| **PQLREKVSKAIDFSQMDLKIDDLSRKGIHTGENPKVVKMKIEPERGAWMSNR** | 121 | 172 | Antigenic | Non-allergen | Non-toxin | Non-homologous |
| **SEVDYIGQFDMRFLNSLAIHEKFDAFMN** | 185 | 212 | Antigenic | Non-allergen | Non-toxin | Non-homologous |
| GGNIPTEFHHY | 260 | 270 | Non-antigenic | Allergen | Non-toxin | Non-homologous |
| **NTNHRHSVLDNTN** | 283 | 295 | Antigenic | Allergen | Non-toxin | Non-homologous |
| **LQDLSLYITETDNAGLKEYKRMEKWTKKS** | 375 | 403 | Antigenic | Allergen | Non-toxin | Non-homologous |

**Table S10.** Primary sequence architecture and modular design of candidate multi-epitope Mpox virus (MPXV) vaccine constructs. Full-length amino acid sequences of engineered chimeric constructs (MVX-1, MVX-2, and MVX-3) illustrating the integration of an N-terminal adjuvant linked via an EAAAK spacer, prioritized MHC Class I epitopes joined by AAY linkers, MHC Class II epitopes joined by GPGPG linkers, and linear B-cell epitopes joined by bi-lysyl (KK) linkers. Total residue lengths are summarized for each candidate.

| Name of the vaccine | Vaccine constructs (AA sequences) | Number of amino acids |
|---|---|---|
| **MVX-1** | EAAAKGIINTLQKYYCRVRGGRCAVLSCLPKEEQIGKCSTRGRKCCRRKKEAA<br>AKSSQDLSFIYAAYYIHPILMALAAYHIDPLGTNVAAYDSDDGFYEWAAYFLA<br>DKKMTLAAYNLQDLSLYIAAYNLLCHIYSLAAYLTTRYHSIYAAYYTDLVISKI<br>AAYSSDELDQKKAAYLPTSIPLAYGPGPGHPILMALFGIKLPALGPGPGYYIHPI<br>LMALFGIKLGPGPGYIHPILMALFGIKLPGPGPGIHPILMALFGIKLPAGPGPGVK<br>SYNYMLLVNRLTEGPGPGTHPLVLIHSRASPRIKKAGLKSSYTNQTLSLVDEDH<br>IHTSDKSKKGARNQRTKPKRTGNDQSQQPIKKDCMVSIDEVASTHDWSTRKKQ<br>NKNMFGGKKDQYISARHIKKSEVDYIGQFDMRFLNSLAIHEKFDAFMNKK | 428 |
| **MVX-2** | EAAAKMAKLSTDELLDAFKEMTLLELSDFVKKFEETFEVTAAAPVAVAAAGA<br>APAGAAVEAAEEQSEFDVILEAAGDKKIGVIKVVREIVSGLGLKEAKDLVDGA<br>PKPLLEKVAKEAADEAKAKLEAAGATVTVKEAAAKSSQDLSFIYAAYYIHPIL<br>MALAAYHIDPLGTNVAAYDSDDGFYEWAAYFLADKKMTLAAYNLQDLSLYI<br>AAYNLLCHIYSLAAYLTTRYHSIYAAYYTDLVISKIAAYSSDELDQKKAAYLPT<br>SIPLAYGPGPGHPILMALFGIKLPALGPGPGYYIHPILMALFGIKLGPGPGYIHPIL<br>MALFGIKLPGPGPGIHPILMALFGIKLPAGPGPGVKSYNYMLLVNRLTEGPGPG<br>THPLVLIHSRASPRIKKAGLKSSYTNQTLSLVDEDHIHTSDKSKKGARNQRTKP<br>KRTGNDQSQQPIKKDCMVSIDEVASTHDWSTRKKQNKNMFGGKKDQYISARH<br>IKKSEVDYIGQFDMRFLNSLAIHEKFDAFMNKK | 513 |
| **MVX-3** | EAAAKMAENPNIDDLPAPLLAALGAADLALATVNDLIANLRERAEETRAETRT<br>RVEERRARLTKFQEDLPEQFIELRDKFTTEELRKAAEGYLEAATNRYNELVERG<br>EAALQRLRSQTAFEDASARAEGYVDQAVELTQEALGTVASQTRAVGERAAKL<br>VGIELPGKAEAAGKKAQKAIAKAPAKKASAKKAPAKKAPAKKAAAKKVTQK<br>EAAAKSSQDLSFIYAAYYIHPILMALAAYHIDPLGTNVAAYDSDDGFYEWAAY<br>FLADKKMTLAAYNLQDLSLYIAAYNLLCHIYSLAAYLTTRYHSIYAAYYTDLVI<br>SKIAAYSSDELDQKKAAYLPTSIPLAYGPGPGHPILMALFGIKLPALGPGPGYYI<br>HPILMALFGIKLGPGPGYIHPILMALFGIKLPGPGPGIHPILMALFGIKLPAGPGPG<br>VKSYNYMLLVNRLTEGPGPGTHPLVLIHSRASPRIKKAGLKSSYTNQTLSLVDE<br>DHIHTSDKSKKGARNQRTKPKRTGNDQSQQPIKKDCMVSIDEVASTHDWSTRK<br>KQNKNMFGGKKDQYISARHIKKSEVDYIGQFDMRFLNSLAIHEKFDAFMNKK | 588 |

**EAAAK (rigid α-helical linker preventing domain interference), AAY (proteasomal cleavage enhancement), GPGPG (prevents junctional epitope formation and facilitates MHC-II antigen presentation), and KK (cathepsin B cleavage site for B-cell presentation).*

**Table S11.** Comparative immunogenic safety and biophysical profiling of candidate multi-epitope Mpox vaccine constructs. Antigenicity scores were determined via VaxiJen (viral model cutoff >0.45) and allergenicity assessed via AllerTOP for constructs MVX-1, MVX-2, and MVX-3. Primary sequence physicochemical properties were modeled via the ExPASy ProtParam server, including total length, molecular weight (MW in Da), theoretical isoelectric point (pI), extinction coefficient ($M^{-1}$ $cm^{-1}$ at 280 nm), aliphatic index (AI), instability index (II), and grand average hydropathicity (GRAVY). All constructs demonstrate thermal stability (AI > 80), stability in solution (II <40.0), and hydrophilic solubility profiles (GRAVY <0).

| Name of the vaccine constructs | Total amino acids | Antigenicity | Allergenicity | Molecular weight (MW) | Theoretical pI | Ext. coefficient (in $M^{-1}$ $cm^{-1}$) | Aliphatic index (AI) | Instability index (II) | Grand average of hydropathicity (GRAVY) |
|---|---|---|---|---|---|---|---|---|---|
| **MVX-1** | 428 | 0.64 | Non-allergen | 47346.02 | 9.49 | 54710 | 86.75 | 36.15 | −0.23 |
| **MVX-2** | 513 | 0.58 | Non-allergen | 55625.30 | 8.40 | 51355 | 92.79 | 31.34 | −0.09 |

| | | | | | | | | | |
|---|---|---|---|---|---|---|---|---|---|
| **MVX-3** | 588 | 0.54 | Non-allergen | 64361.01 | 9.34 | 55825 | 86.68 | 37.14 | −0.32 |

**Table S12.** Secondary structure composition of engineered Mpox vaccine constructs evaluated via SOPMA. Proportional distribution of conformational elements, including *α*-helices, *β*-sheets (extended strands), and random coil conformations across the candidate multi-epitope constructs (MVX-1, MVX-2, and MVX-3). The prioritized candidate MVX-3 is enriched in ordered α-helical content (59.35%) alongside a compacted random coil framework (28.91%), conferring structural stability and fold integrity.

| Name of the vaccines | α-helix percentage of amino acids | β-sheet percentage of amino acids | Coil structure percentage of amino acids |
|---|---|---|---|
| **MVX-1** | 17.29% | 16.36% | 66.36% |
| **MVX-2** | 42.30% | 13.65% | 44.05% |
| **MVX-3** | 59.35% | 11.73% | 28.91% |

**Table S13.** Molecular docking benchmarking and binding free energy estimation of engineered Mpox vaccine constructs complexed with human Toll-like receptors (TLRs). Macromolecular docking scores across human immune receptors TLR1 (PDB ID: 6NIH), TLR2 (PDB ID: 3A7C), TLR3 (PDB ID: 2A0Z), and TLR4 (PDB ID: 4G8A) were evaluated via ClusPro, HDOCK, and HawkDock. Binding free energies ($\Delta G_{bind}$ in kcal mol$^{-1}$) were estimated via the MM-GBSA module of HawkDock. Receptors exceeding the maximum sequence length parameter (>1,000 amino acids) of the HawkDock algorithm are denoted as NA. Prioritized receptor–construct complexes are shown in bold and selected for subsequent all-atom molecular dynamics validation.

| Name of the vaccines | Name of the targets | PDB IDs of the targets | ClusPro energy score | HDock docking score | HawkDock docking score | MM-GBSA (binding free energy, in kcal mol$^{-1}$) |
|---|---|---|---|---|---|---|
| | TLR1 | 6nih | −1024.1 | −348.19 | NA | NA |
| | TLR2 | 3a7c | −1136.8 | −304.60 | −5313.12 | −24.42 |
| | TLR3 | 2a0z | −1131.9 | −361.85 | −5320.86 | 41.81 |
| MVX-1 | TLR4 | 4g8a | −1086.4 | −337.30 | NA | NA |
| | TLR8 | 3w3m | −1166.2 | −342.87 | −6123.01 | −27.11 |
| | TLR9 | 4qdh | −1488.4 | −326.09 | −5846.97 | −23.06 |
| | 3C5J | 3c5j | −1135.2 | −286.56 | −5485.54 | −47.5 |
| | TLR1 | 6nih | −1140.5 | −340.00 | NA | NA |
| | TLR2 | 3a7c | −1126.3 | −330.99 | −3001.76 | 0.89 |
| | TLR3 | 2a0z | −1138.6 | −369.58 | −2999.92 | −1.56 |
| | TLR4 | 4g8a | −1243.9 | −380.78 | NA | NA |
| MVX-2 | TLR8 | 3w3m | −1263.4 | −373.24 | −3157.69 | −1.22 |
| | TLR9 | 4qdh | −1374.7 | −364.58 | −3026.68 | 2.38 |
| | 3C5J | 3c5j | −1240.2 | −291.60 | −2598.52 | −2.87 |
| | TLR1 | 6nih | −1129.0 | −348.19 | NA | NA |
| | TLR2 | 3a7c | −1056.8 | −304.60 | −3654.34 | −16.26 |
| **MVX-3** | **TLR3** | 2a0z | −1099.4 | −361.85 | −2379.80 | −16.33 |
| | TLR4 | 4g8a | −1135.0 | −322.67 | −2833.38 | −5.09 |
| | TLR8 | 3w3m | −1159.1 | −342.87 | −3213.07 | −25.6 |

| | | | | | |
|---|---|---|---|---|---|
| | TLR9 | 4qdh | −1035.3 | −326.09 | −2634.16 | −6.62 |
| | 3C5J | 3c5j | −1169.6 | −286.56 | −2436.33 | −19.36 |

**Table S14.** *In silico* engineered full-length mRNA transcript sequence encoding the prioritized Mpox vaccine candidate (MVX-3). Primary sequence architecture of the 1,780-nucleotide optimized transcript, incorporating the 5' untranslated leader initiation window, multi-epitope open reading frame, and translation termination elements generated via the RiboDecode generative recurrent neural network framework. Thermodynamic stability profiling confirms a global minimum free energy (MFE) of ΔG = −678.1 kcal $mol^{-1}$ with an unconstrained, accessible 5' initiation portal (ΔG = −11.6 kcal $mol^{-1}$).

| Construct | Sequence | Transcript length (nt) |
|---|---|---|
| **MVX-3** | AAGGAGATATACCGAAGCGGCGGCGAAAATGGCGGAAAACCCGAACATTGATGATCTGCCGGCGC<br>CGCTGCTGGCGGCGCTGGGCGCGGCGGATCTGGCGCTGGCGACCGTGAACGATCTGATTGCGAAC<br>CTGCGTGAACGTGCGGAAGAAACCCGTGCGGAAACCCGTACCCGTGTGGAAGAACGTCGTGCGCG<br>TCTGACCAAATTTCAGGAAGATCTGCCGGAACAGTTTATTGAACTGCGTGATAAATTTACCACCGA<br>AGAACTGCGTAAAGCGGCGGAAGGCTATCTGGAAGCGGCGACCAACCGTTATAACGAACTGGTGG<br>AACGTGGCGAAGCGGCGCTGCAGCGTCTGCGTAGCCAGACCGCGTTTGAAGATGCGAGCGCGCGT<br>GCGGAAGGCTATGTGGATCAGGCGGTGGAACTGACCCAGGAAGCGCTGGGCACCGTGGCGAGCCA<br>GACCCGTGCGGTGGGCGAACGTGCGGCGAAACTGGTGGGCATTGAACTGCCGGGCAAAGCGGAAG<br>CGGCGGGCAAAAAAGCGCAGAAAGCGATTGCGAAAGCGCCGGCGAAAAAAGCGAGCGCGAAAAA<br>AGCGCCGGCGAAAAAAGCGCCGGCGAAAAAAGCGGCGGCGAAAAAAGTGACCCAGAAAGAAGCG<br>GCGGCGAAAAGCAGCCAGGATCTGAGCTTTATTTATGCGGCGTATTATATTCATCCGATTCTGATG<br>GCGCTGGCGGCGTATCATATTGATCCGCTGGGCACCAACGTGGCGGCGTATGATAGCGATGATGGC<br>TTTTATGAATGGGCGGCGTATTTTCTGGCGGATAAAAAAATGACCCTGGCGGCGTATAACCTGCAG<br>GATCTGAGCCTGTATATTGCGGCGTATAACCTGCTGTGCCATATTTATAGCCTGGCGGCGTATCTGA<br>CCACCCGTTATCATAGCATTTATGCGGCGTATTATACCGATCTGGTGATTAGCAAAATTGCGGCGT<br>ATAGCAGCGATGAACTGGATCAGAAAAAAGCGGCGTATCTGCCGACCAGCATTCCGCTGGCGTAT<br>GGCCCGGGCCCGGGCCATCCGATTCTGATGGCGCTGTTTGGCATTAAACTGCCGGCGCTGGGCCCG<br>GGCCCGGGCTATTATATTCATCCGATTCTGATGGCGCTGTTTGGCATTAAACTGGGCCCGGGCCCG<br>GGCTATATTCATCCGATTCTGATGGCGCTGTTTGGCATTAAACTGCCGGGCCCGGGCCCGGGCATT<br>CATCCGATTCTGATGGCGCTGTTTGGCATTAAACTGCCGGCGGGCCCGGGCCCGGGCGTGAAAAGC<br>TATAACTATATGCTGCTGGTGAACCGTCTGACCGAAGGCCCGGGCCCGGGCACCCATCCGCTGGTG<br>CTGATTCATAGCCGTGCGAGCCCGCGTATTAAAAAAGCGGGCCTGAAAAGCAGCTATACCAACCA<br>GACCCTGAGCCTGGTGGATGAAGATCATATTCATACCAGCGATAAAAGCAAAAAAGGCGCGCGTA<br>ACCAGCGTACCAAACCGAAACGTACCGGCAACGATCAGAGCCAGCAGCCGATTAAAAAAGATTGC<br>ATGGTGAGCATTGATGAAGTGGCGAGCACCCATGATTGGAGCACCCGTAAAAAACAGAACAAAAA<br>CATGTTTGGCGGCAAAAAAGATCAGTATATTAGCGCGCGTCATATTAAAAAAAGCGAAGTGGATT<br>ATATTGGCCAGTTTGATATGCGTTTTCTGAACAGCCTGGCGATTCATGAAAAATTTGATGCGTTTAT<br>GAACAAAAAATAA | 1780 |

### Supplementary Figures

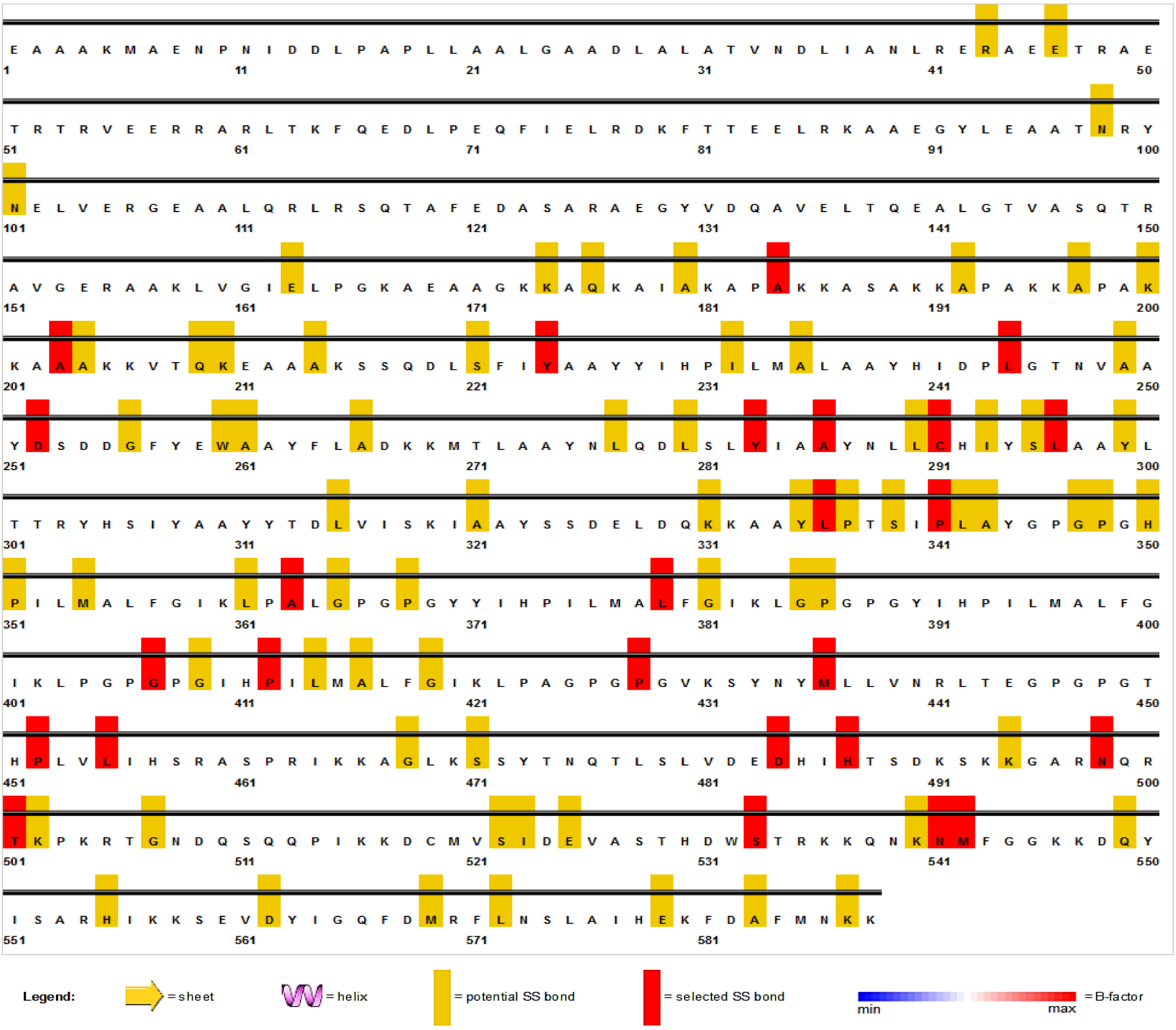


**Figure S1**. Two-dimensional mapping of disulfide bridge engineering and mutable residue profiles for conformational stabilization of the MVX-3 vaccine construct. Prediction made using the Disulfide by Design v2.0 (DbD2) server on the tertiary model of the multi-epitope construct. The upper yellow blocks indicate pairs of residues with steric potential favorable for the formation of disulfide bonds ($\chi_3$ torsional angle and optimal interatomic distance), while the solid red blocks highlight the pairs of mutated/selected cysteines (final SS bonds) introduced to restrict conformationally labile regions without perturbing epitopic determinants.

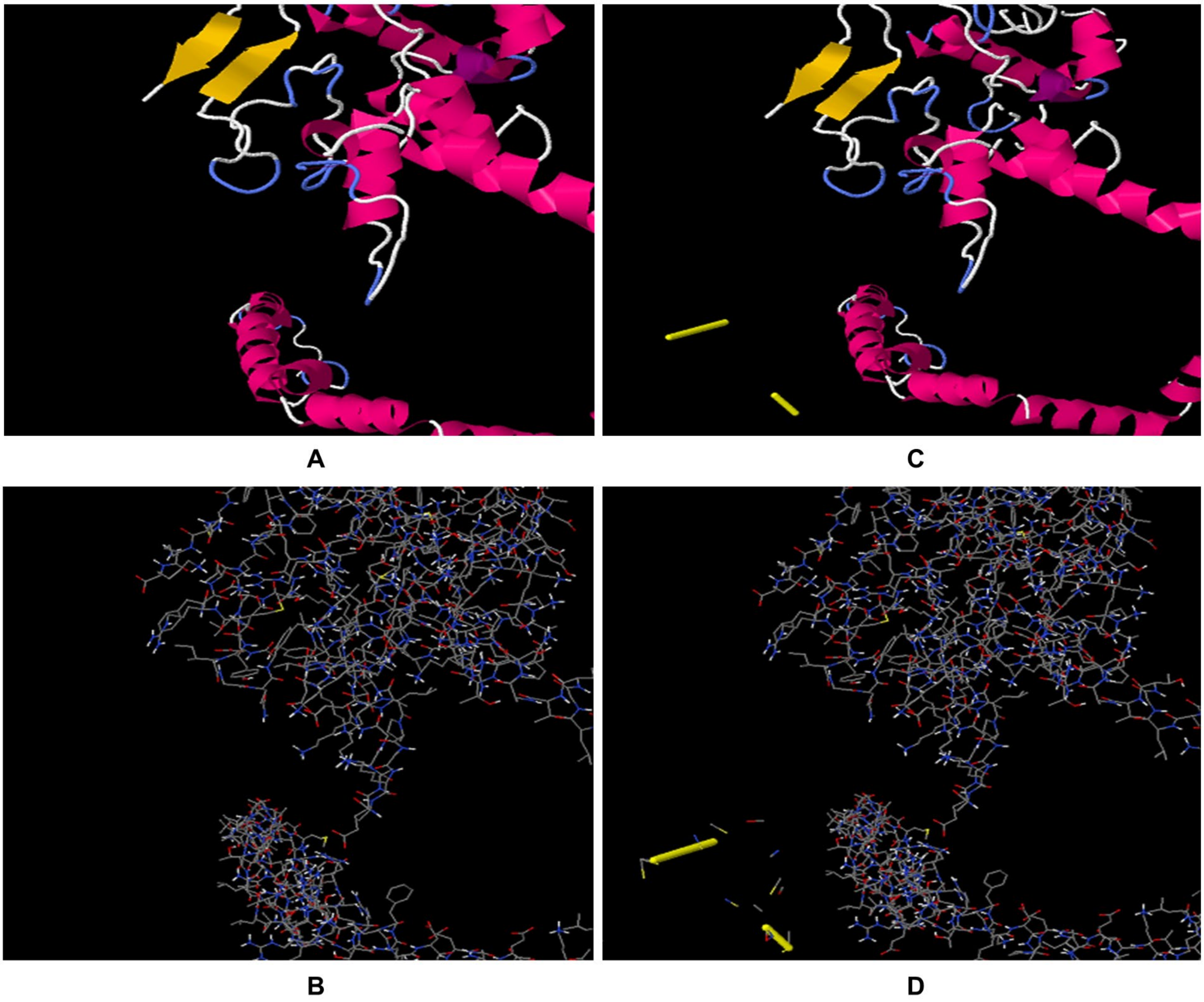


**Figure S2.** Comparative three-dimensional structural architecture of the MVX-3 vaccine constructed before and after *in silico* disulfide engineering. Structural modeling generated via Disulfide by Design v2.0 (DbD2). (A) Cartoon ribbon and (B) all-atom wireframe representation of the wild-type (unmutated) MVX-3 tertiary model. (C) Cartoon ribbon and (D) corresponding wireframe rendering of the engineered mutant construct, illustrating the spatial introduction of stereochemically optimized disulfide bridges (highlighted as solid yellow cylinders) engineered to enhance conformational rigidity without disrupting antigenic epitope display.
